# Population responses in V1 encodes stimulus visibility in backward masking

**DOI:** 10.1101/2025.03.30.642841

**Authors:** Itiel Dennis, Hadar Edelman-Klapper, Gal Gadi Raveh, Osnat Bar-Shira, Shany Nivinsky Margalit, Hamutal Slovin

## Abstract

The visibility of a briefly presented stimulus is diminished when followed by a mask, a phenomenon known as backward masking (BM). As the interval between the stimulus and the mask (stimulus-to-mask onset asynchrony, SOA) becomes shorter, the stimulus visibility decreases. Yet, the neural mechanisms underlying BM remains poorly understood. To investigate this, monkeys were trained to discriminate oriented targets in pattern BM. Using Voltage-sensitive dye imaging we measured the population responses in the primary visual cortex (V1). The behavioral performance in short SOAs was lower and reaction times were slower. Population response in V1 showed a figure-ground modulation (FGm) with lower values for short SOAs. A classification analysis revealed the differences in temporal dynamics of the model accuracy in relation to stimulus visibility and spatial rearrangements of the informative neural activity. These results suggest that the mask interferes with FGm in V1 and possibly leads to diminished stimulus visibility.

## Introduction

The paradigm of visual masking has been used for over a century to disrupt visual perception by presenting a masking stimulus that is in close proximity in time and space to a target stimulus (Kahneman, 1968; Enns and Lollo, 2000; Fahrenfort et al., 2007; Macknik and Martinez 2007; Bachmann and Francis, 2014). Here we used the backward masking (BM) paradigm, in which the visibility of a stimulus is suppressed by a subsequent masking stimulus (Karni & Sagi, 1991; Breitmeyer and Ogmen, 2000). This effect is not simply attributed to the brief presentation of the first stimulus, because when it is not followed by a mask, the stimulus can be clearly seen. Behavioral studies reported that stimulus visibility in BM depends on the time interval between the stimulus onset and the mask onset (stimulus-to-mask onset asynchrony; SOA) such as the SOA becomes shorter, the stimulus visibility decreases (Breitmeyer and Ganz, 1976; Karni and Sagi, 1991, 1993; Francis et al., 2004). The effects of BM have been linked to the neural activity in early visual cortex (Karni and Sagi, 1991; Macknick & Livingstone, 1998; Breitmeyer & Ogmen, 2000; Lamme et al., 2002; Huang et al., 2006; Fahrenfort et al., 2007; Alwis et al. 2016; Gale et al., 2024). However, despite the extensive research, the neuronal mechanisms in the visual cortex underlying BM are still not well understood.

Several mechanisms were suggested to explain the masking effects: including temporal integration of sensory signals causing target-mask merging (Kinsbourne and Warrington 1962; Kahneman et al., 1968; Coltheart and Arthur, 1972; Felsten and Wasserman, 1980; Gale et al. 2024), lateral inhibition in the feedforward visual pathway (Francis, 1997; Macknik & Livingstone, 1998; Macknik and Martinez-Conde 2007; Huang et al., 2006) and disruption of cortical feedback (Bridgeman, 1980; Di Lollo et al., 2000; Ro et al., 2003; Fahrenfort et al., 2007). Lamme et al. (2002) extended this theory to suggest that the mask interrupts the reentrant processing from higher visual areas back to V1 of monkeys i.e. that the masking interferes with the figure-from-ground processing resulting in decreased stimulus visibility (Lamme et al., 2002). Yet, how the figure-ground processing is interrupted in V1 and the relation to behavioral performance ans stimulus visibility in BM is poorly understood To investigate this question, we used Voltage-sensitive dye imaging (VSDI) to measure the population responses in the primary visual cortex (V1) of monkeys trained on pattern BM. We found that the animal’s behavioral performance decreased in short SOAs and reaction times became slower relative to long SOAs. The V1 population response showed a figure-ground modulation (FGm) where the FGm in late time (∼200 ms after stimulus onset) was higher in longer SOAs where stimulus visibility was better. A classification analysis revealed spatial reorganization of the informative regions in V1 that could discriminate between trials with high and low visibility. Our results suggest that the mask interferes with figure-ground segregation in V1, possibly leading to diminished stimulus visibility.

## Results

### Behavioral performance in the backward masking task

Two monkeys were trained on a texture discrimination task (TDT) with pattern backward masking (BM, Fig. 1). The monkeys were required to discriminate between two texture stimuli that included either a horizontal or a vertical target (Fig. 1Ai). The texture stimuli were comprised of oriented line segments, while a target consisted of three horizontally or vertically aligned lines which were segmented from the background lines only by their orientation difference (Fig. 1Ai). On each trial, the monkeys acquired and then maintained fixation on a small fixation point, for a random time interval (Fig. 1Aii). Then, a texture stimulus, either a horizontal or vertical stimulus, was turned on the screen. The stimulus appeared for a brief time (40 ms) and after a short period of time a mask stimulus was turned on for 100 ms (Fig. 1Ai, right). The time interval between the stimulus onset and mask onset is termed as the stimulus-to-mask onset asynchrony (SOA), and its duration varied (50, 75, 100 &150 ms) across the different recording sessions. Following the mask offset, the monkeys were required to report the target orientation by making a saccade (Fig. 1Aii).

**Figure 1:**
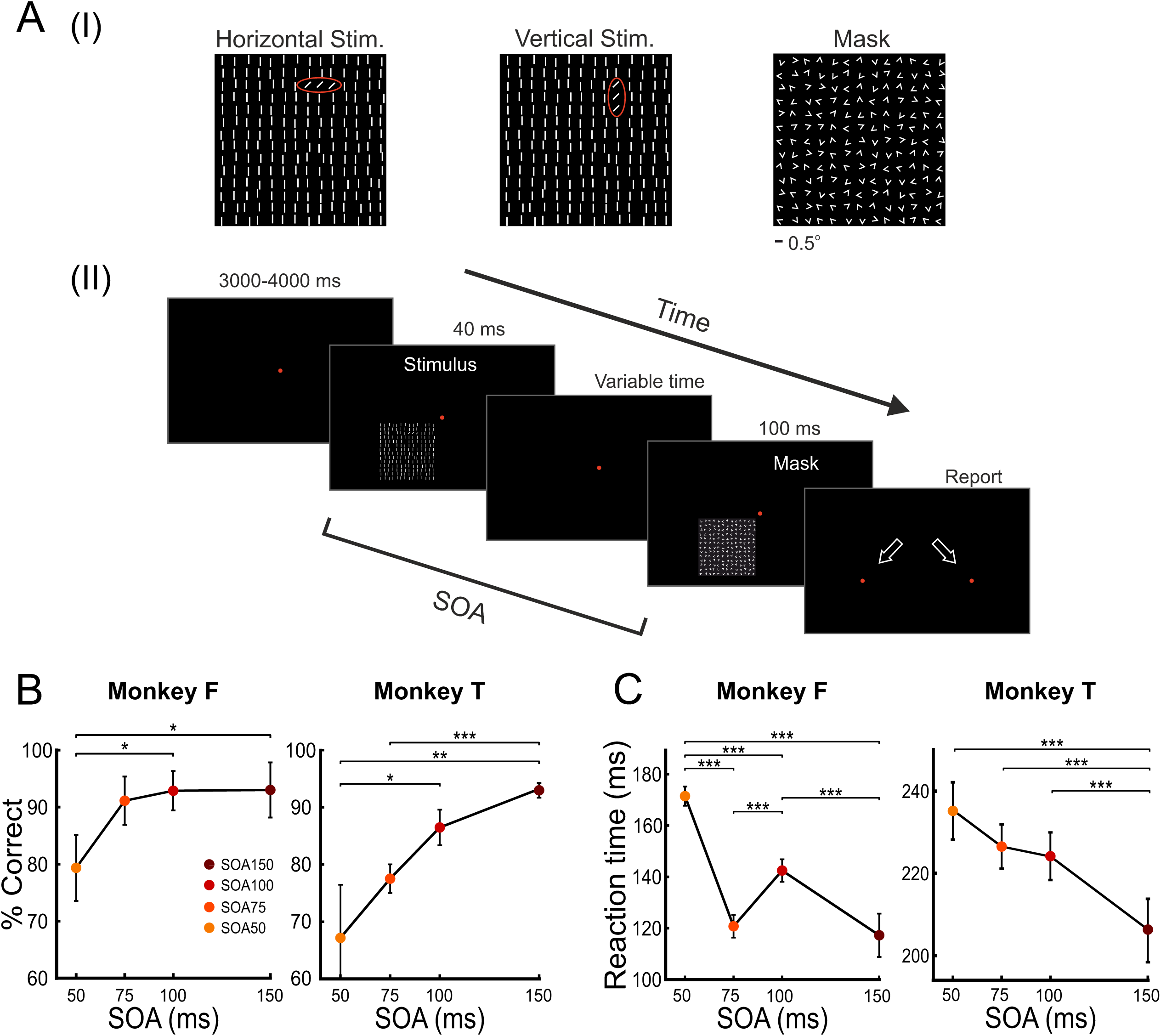
T**e**xture **discrimination with backward masking paradigm and behavioral performance. A.** The stimuli and mask. **(i)** The texture stimuli were composed of three lines of bars oriented horizontally (left: horizontal stimulus) or vertically (middle: vertical stimulus), embedded in background of oriented bars (see Methods). The red ellipses depict the target’s elements: horizontal target (left) and vertical target (middle). The red ellipses are for illustration only (not visible to the monkey). Right: The mask stimulus, comprised of random oriented V-shape elements. **(ii)** The Texture discrimination with backward masking (BM) paradigm (see Methods). The monkey had to fixate over a small fixation point (red point) for a random interval of 3000-4000 ms until the stimulus was turned on for 40 ms. After a variable period of time that is defined as stimulus-to-mask onset asynchrony (SOA), a mask was tuned on for 100 ms. The SOA duration values varied between sessions: 150, 100, 75 and 50 ms. Following the mask and fixation point offset, two small lateral points appeared on the screen and the monkey had to report what was the target’s orientation (horizontal or vertical) by performing a saccade to the left or to the right point. **B.** The behavioral performance in the BM task was evaluated by computing the percentage of correct trials (fraction of correct trials out of total number of trials) as function of SOA, for monkeys F and T, across sessions. Numbers are mean ± SEM across sessions. Monkey F: n = 8, 6, 8 & 6 sessions for SOAs 50, 75, 100 & 150 ms respectively. Monkey T: n = 6, 12, 10 & 12 sessions for SOAs 50, 75, 100 & 150 ms respectively. **C.** Reaction time (RT) as function of SOA for monkeys F and T. The RT for each SOA, was averaged across trials. Monkey F: n = 173, 111, 171, 106 trials for SOAs 50, 75, 100 & 150 respectively. Monkey T: n = 124, 354, 358, 399 trials for SOAs 50, 75, 100 & 150. * p<0.05 ** p < 0.01 *** p< 0.001, rank-sum test between sessions for behavioral performance and between trials for RT.

Next, we assessed the monkeys’ behavioral performance in the BM task and found that it depended on the SOA duration, as was previously reported. The detection rate of horizontal and vertical stimuli (defined as: the percentage of correct trials out of the total number of trials, see Methods) was higher at long SOA (150 ms) and significantly decreased at short SOA (50 ms; Fig. 1B). In Monkey F, the percentage of correct trials was 93.01 ± 4.84% (mean±sem) in the longest SOA and it decreased to 79.35 ± 5.80% in the shortest SOA (p < 0.001; Wilcoxon rank-sum test). In monkey T, the percentage of correct trials was 92.47 ± 1.41% in the longest SOA and it decreased to 67.44 ± 8.05 % (p < 0.001). Next, we computed the reaction time (RT; defined as the time interval from the mask offset until the reporting saccade has landed) of the animals, which revealed an inverse trend: trials with longer SOA had shorter RT whereas trials with shorter SOA had a longer RT (Fig. 1C). In Monkey F the RT increased from 117.25 ± 8.44 ms in long SOA to 171.47 ± 3.74 ms in short SOA. In monkey T the RT increased from 207.28 ± 7.45 ms in long SOA to 234.45 ± 7.34 ms in short SOA (P < 0.001 for both monkeys). The behavioral results are consistent with previous reports from humans (Karni and Sagi, 1991; Censor et al., 2009; Schwartz et al., 2002; Scoups and Orban, 1996; Yotsumoto et al., 2008; Bacon-Mace et al. 2005) and monkeys (Lamme et al. 2002; Op de Beeck et al. 2007; Rolls 2004; Thompson & Schall 1999) that performed a BM task. Moreover, the RTs in error trials (trials in which the animal has reported on perceiving the opposite target to that presented) were longer than the RTs in the correct trials (Supplementary Fig. S1A, S1B). The RT in error trials were significantly longer by an average of 57.6 ms (p < 0.001 for all SOAs combined) and 38.7 ms (p < 0.001 for all SOAs combined) in monkey F and T respectively. Moreover, when dividing the RTs into bins according to their lengths, we found that trials with faster RTs were associated with higher fraction of correct trials within each SOA (Supplementary Fig. S1C, S1D). The longer RT in error trials, as well as the dependency of the percent correct on the RT length, support the notion that trials with longer RT reflect poorer visibility of the target.

While the animals performed the BM task, we recorded the population response in area V1 at high spatial (mesoscale, 170^2^ μm^2^/pixel) and temporal resolution (100Hz) using voltage-sensitive dye imaging (VSDI; Slovin et al. 2002). The fluorescence dye signal of each pixel reflects the sum of membrane potential from a population of neurons (rather than single cells) emphasizing subthreshold membrane potentials but also reflecting also suprathreshold membrane potentials (i.e. spiking activity of neuronal populations; Jancke et al., 2004; Ayzenshtat et al., 2010).

### Mapping the targets and background parts of the stimulus onto V1

To study the population responses in the BM task, we first needed to map the different stimulus’ parts (target and background) onto the V1 imaging area (see Methods). Therefore, on each recording day, the monkeys initially performed a simple fixation paradigm, where a vertical target alone (Fig. 2Ai) or a horizontal target alone (Fig. 2Bi), without the background elements, were briefly presented. The two targets shared one common element (Fig. 2Ci, yellow element), while the other elements did not overlap (Fig. 2Ci, red and blue elements for the vertical and horizontal target, respectively). Next, we used VSDI to obtain the retinotopic mapping, i.e. the spatial pattern of the population response in V1 to each of the targets. Figure 2Aii and 2Bii shows the population response map for the vertical target and horizontal target at peak response amplitude (Fig. 2Di, 2Dii shows the temporal evolvement of the distinct activation patterns to each of the targets in V1). Based on the retinotopic activation maps we defined the following regions of interest (ROIs) in V1: (1) Vertical ROI and (2) Horizontal ROI that were defined by fitting an elliptical contour to the activation patches in V1 evoked by the vertical or horizontal targets (Fig. 2Aii, 2Bii, black contours). Next, we defined a common-ROI, corresponding to the common element between the two targets: the overlapped V1 region between the vertical ROI and the horizontal ROI (Fig. 2Cii, yellow contour). The remaining non-overlapping parts between the vertical and horizontal ROIs were defined as vertical-only and horizontal-only ROIs (Fig. 2Cii, red and blue contours, corresponding to the non-overlapping elements in the two targets). The V1 area outside the target ROIs was defined as the general background region (Fig. 2Cii).

**Figure 2.**
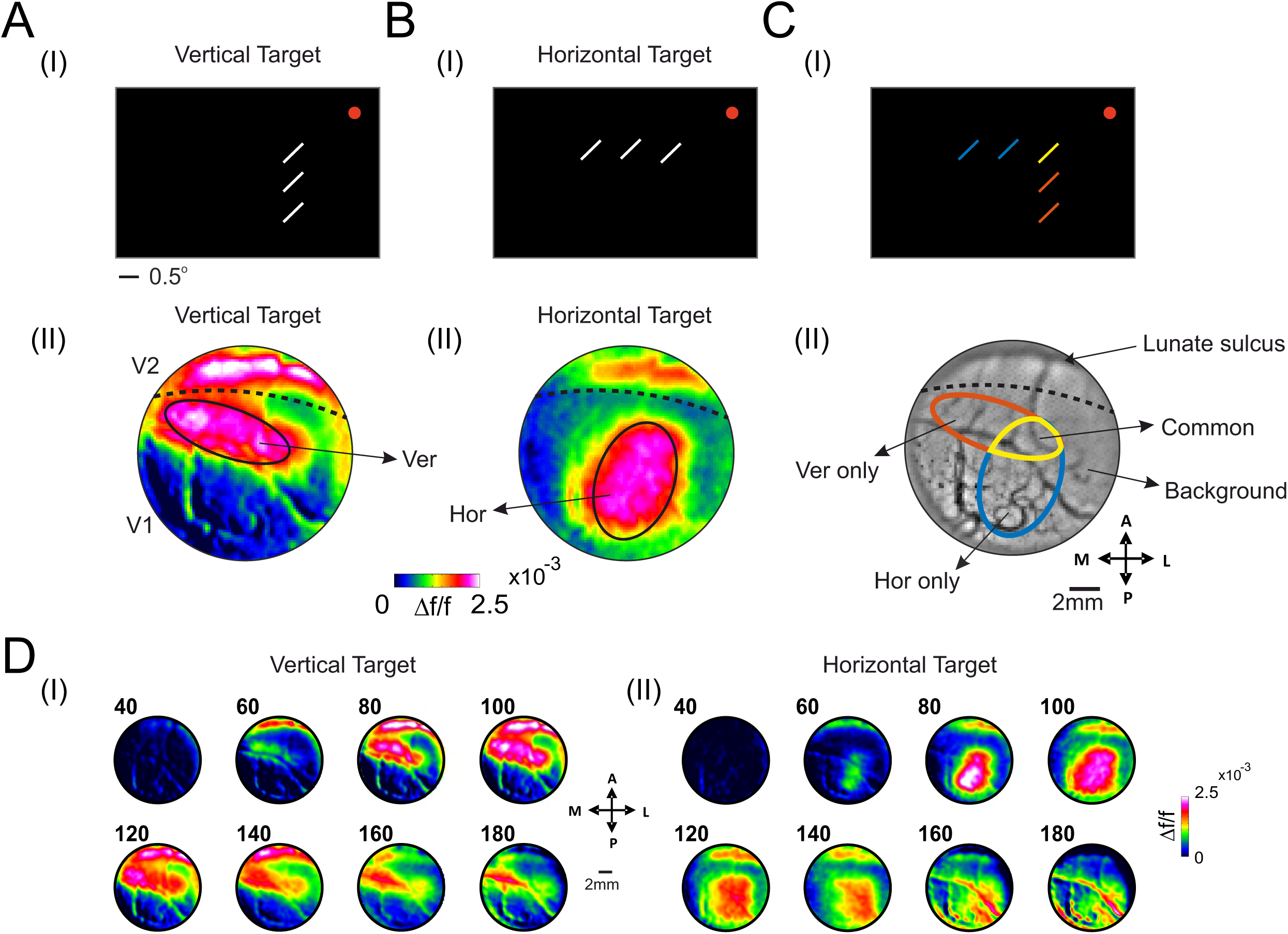
: Mapping the targets and background regions onto the retinotopic map of V1. A. (i) Illustration of the vertical target, without background elements. Fixation point is marked by a red point. (ii) The VSD maps at peak response (mean over 70-110 ms after stimulus onset, n=11 trials, example session monkey F) for the vertical target, maps are color coded. The black contour represents the vertical (Ver) ROI. The dashed line represents the border between V1 and V2. B. Same as in A but for the horizontal target and horizontal (Hor) ROI (n=8 trials, example session monkey F). C. The vertical and horizontal targets share a common element on the display screen, and therefore also over the retinotopic map in V1. (i) The blue elements represent the bar elements belonging to the horizontal target only, red elements represent the bars belonging to the vertical target only and yellow bar denotes the common element of both targets. (ii) Same ROIs as in Aii and Bii superimposed over the blood vessels map from the same recording session. The red contour represents the vertical-only ROI (Ver only), the blue contour represents the horizontal-only ROI (Hor only) and the yellow contour represents the common ROI that is corresponding to the common element of the two targets. D. A sequence of VSD maps showing the spatio-temporal development of population responses to the vertical target (i) or horizontal target (ii). Numbers above the maps indicate time (in ms) after stimulus onset, same data as in A(ii) and B(ii). Abbreviations: A- Anterior; P-Posterior; L- Lateral; M- Medial.

### The V1 response in the backward masking paradigm and the relation to the stimulus and mask alone evoked responses

To investigate the neural response for the stimulus from that of the mask, the monkeys performed a fixation paradigm while the stimulus alone (40 ms duration; not followed by a mask; Fig. 3Ai) was presented. In a different set of fixation trials, the mask alone appeared (100 ms duration, as in the BM task; Fig. 3Bi; see Methods). Figure 3Aii and Fig. 3Bii show the temporal evolvement of the VSD maps following the stimulus onset or mask onset (the time points of the maps are depicted on the time courses (TCs) of the V1 VSD signal in Fig. 3Aiii, 3Biii). Unlike the target alone stimuli, which elicited distinct patches of activation within the V1 region (Fig. 2), the full texture stimulus and mask stimulus elicited a wide-spread activation across the entire imaged V1 area. Within this wide-spread activation (that is expected due the multiple oriented bars and stimulus size) the activity evoked by the target elements could not be discriminated easily from the population activity evoked by the background elements. Next, we computed the VSD TC (mean response over all V1 pixels and across trials) for the stimulus alone condition (Figure 3Aiii) and for the mask alone condition (Figure 3Biii; the TC of the evoked population response is longer than the stimulus duration itself, which is typical for V1 response in studies using very brief stimuli e.g. Lamme at al. 2002; in addition, the stimuli were flashed over a dark background and therefore the population response decay slowly to baseline level within the time window, see Methods).

**Figure 3.**
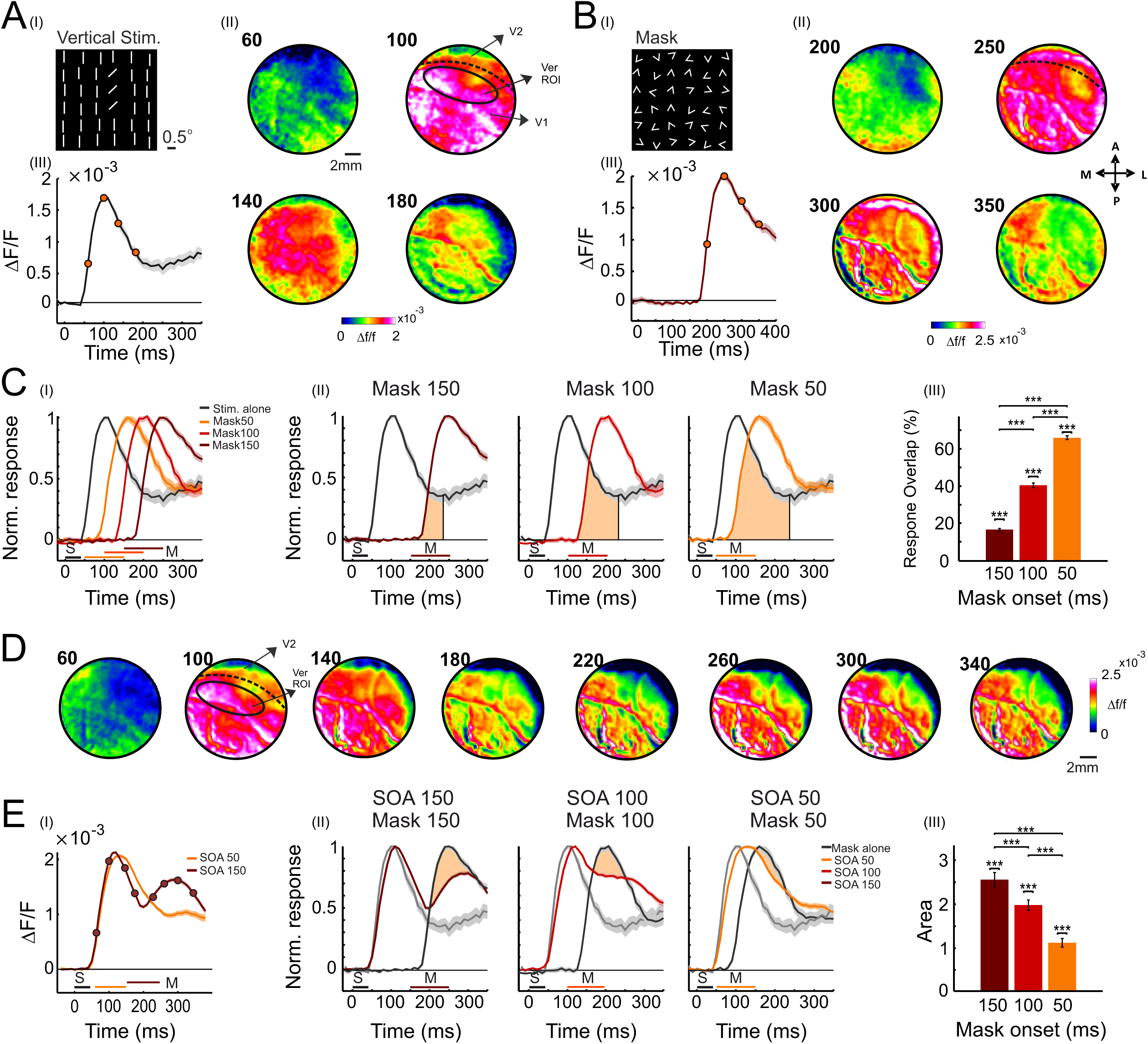
A widespread VSD response in V1 in the BM task and the relation to the stimulus alone and mask alone responses. **A.** The VSD response to the stimulus alone. **(i)** The vertical target and background elements from a vertical stimulus (only part of the stimulus image is shown). **(ii)** The temporal evolvement of the VSD maps following the stimulus onset, an example session (n=5 trials, monkey F). **(iii)** The VSD TC (mean over V1 pixels), same data as in Aii. The red dots on the VSD TC correspond to the times of VSD maps in A (ii). **B.** Same as in A, but for the mask alone stimulus (only part of the mask image is shown), an example session (n=9 trials). **C (i)** The normalized TC of the VSD signal for the stimulus alone condition (black line, same data as in Aii) or Mask alone conditions (brown traces). All TCs are normalized to peak response and are arranged for increasing onset times of the Mask alone stimulus: 50, 100, 150 ms after stimulus onsets (these timing matches the 50, 100 and 150 SOAs in the BM task). n=18, 16, 15 trials accordingly. **(ii)** The area of overlap between the stimulus (alone) and the mask (alone) evoked response computed separately for each mask onset time (see Methods). **(iii)** The grand analysis of the area of overlap between stimulus alone and mask alone responses. Data is from monkey F (n=33 trials for stimulus alone; n=15, 26, 29 trials for mask alone, at 150, 100 and 50 ms onset from stimulus onset). **D.** A sequence of VSD maps in the BM task, SOA 150 ms, example session (monkey F, n=18 trials). The maps show a biphasic response: the first peak population response corresponds to the stimulus onset and the second peak population responses corresponds to the mask onset. The numbers above the maps denote the time (in ms) after stimulus onset, maps are separated by 40 ms. **E. (i)** The TC of the population response for the example session in D (n=36 trials, SOA 150 ms) and for SOA 50 (n=35 trials). The VSD responses at the first peak are not significantly different, whereas the VSD response at the second peak (280-300 ms) are significantly different (p <<0.001). **(ii).** The normalized TC of the VSD signal (mean over V1 pixels) in the stimulus alone condition (gray curve) or Mask alone condition (black curve). The TCs are arranged for the different mask onset times (from left to right): 150, 100 and 50 ms (n=15,16,18 trials accordingly). On each panel, a normalized TC for the BM paradigm is shown, with the matched SOA duration (SOA 50: n = 35; SOA 100: n = 36; SOA 150: n = 36). Colored shaded area around the curves represents ±1 SEM over trials. The bars at the bottom of the TCs represent the stimulus duration (S) or the mask (M) in all figures. The TC is averaged over trials and is normalized to peak response of each TC (stimulus alone: n=5). Beige area refers to the difference between the VSD response in the mask only condition and the response to the mask in the BM paradigm. **(iii)** Grand analysis and quantification of the areal difference described in Eii (n = 15, 26 & 29 for SOA150, 100 & 50 respectively in the BM condition; for the mask alone condition see Ciii). * p<0.05; ** p<0.01; *** p<0.001 here and in all figures.

Figure 3Ci shows the TC of the VSD response (mean across trials) to the stimulus alone (black curve) and to the mask alone (orange to brown curves). The onset time of the mask alone was varied to match the mask onset in the different SOAs, from short to long SOAs. The VSD TC is normalized to the peak response of each condition. The results in Fig. 3Ci shows that as the mask onset is earlier in time, the mask alone evoked response appears closer to the stimulus alone response. Figure 3Cii shows the TCs of the VSD signal for the stimulus alone and the mask alone separated for the different onset times of the mask. While for the late mask onset (Mask 150 ms) there is only a small overlap between the stimulus evoked response and the mask evoked response (light orange area, Fig. 3Cii), as the onset time of the mask becomes earlier, the mask evoked response is shifted earlier in time and there is a larger overlap with the stimulus evoked response. For mask 50 (which is equivalent to SOA 50) there is a large overlap between the mask evoked response and the stimulus evoked response (Fig. 3Cii). To quantify the overlap between the response to the stimulus alone and the response to the mask alone, we computed the enclosed area between the two responses, divided by the response to the stimulus alone, for each SOA (Fig. 3Ciii; see Methods). At the longest mask onset (equivalent to SOA150 ms), the overlap area covers 16.5 ± 0.62% of the stimuli-related area, as opposed to a 40.46 ± 1.23% overlap in SOA 100 ms and 66.0 ± 1.02% in SOA 150 ms (p value range: p < 0.01 to p< 0.001, signed-rank for difference from 0 and rank-sum for difference between mask onsets time). These results suggest that as the SOA shortens, the neural response to the mask becomes more disruptive due to a larger overlap with the stimulus-related response. Yet it appears that the early part of the stimulus evoked response (< 100 ms) is not affected by the mask.

Next, we moved on to investigate the V1 response in the BM task, which elicited a wide- spread activation throughout the whole V1 imaging area. The sequence of the VSD maps in Fig. 3D shows the spatio-temporal activation patterns in V1 evoked by a vertical stimulus that is followed by a mask (SOA, 150 ms), in an example session. The VSD maps show two temporal phases of increased neural activation, corresponding to the population response evoked by the stimulus (response window: ∼80-160 ms) and by the mask (response window: ∼220-340 ms). Next, we computed the TC of the population response in the SOA 150 & 50 ms for the same example session in Fig. 3D. Figure 3Ei shows the TCs of the population response in the two SOAs which exhibit an early peak of activation ∼110-130 ms after the stimulus onset with similar amplitude (2.10×10^-3^ ± 2.3×10^-5^ mean±sem across trials, and 2.0×10^-3^ ± 2.4×10^-5^ respectively; non-significant difference between the SOAs; p << 0.001, for difference from zero for both SOAs). Interestingly, the response for the SOA 150 has a second peak of activation later on (∼280-300 ms, 1.60×10^-3^ ± 3.7×10^-5^; p << 0.001 for difference from zero) evoked by the mask onset (150 ms after stimulus onset). However, for the SOA 50 (i.e. mask onset is 50 ms after stimulus onset) there is no clear second phase of increased activation that corresponds to the mask onset. The VSD TC suggests that the response to the mask at SOA 50 was merged with the stimulus evoked activity. The population response for SOA 50, at t=280-300 ms was 9.97×10^-4^ ± 4.6×10^-5^ which is significantly lower from the SOA 150 (p < 0.001) and significantly higher from zero (p < 0.001; signed-rank test).

Next, we investigated the VSD response in the BM in relation to the stimulus alone and mask alone responses. Figure 3Eii shows the population response TC (mean across trials) from the BM condition along with the population response to the stimulus alone and mask alone at matching times to the SOAs. In the VSD response of the BM, the initial response fits with the stimulus alone response, for all SOAs (black curve vs. brown-orange curves). For SOA 150 ms, the second phase of increased activation clearly fit on the time axis, with the response to the mask alone at the matched time. Yet the population response in the BM task, at the time corresponding to the mask, shows a lower response of activation than the response evoked by the mask alone. We termed this phenomenon as mask suppression and to quantify it, we computed the area under the population response to the mask alone condition relative to the population response in the BM task (see Methods). This was computed for the different SOAs (150, 100 & 50; beige area). This quantification is shown in Fig. 3Eiii, revealing that the mask suppression becomes smaller as the SOA becomes shorter (2.56 ± 0.16, 1.98 ± 0.17, 1.12 ± 0.09 for SOAs 150, 100, 50 respectively; p < 0.01 to p <0.001, for difference from 0 and for difference between SOAs).

In summary, it appears that the early part of the stimulus evoked response (< 100 ms) is not affected by the mask at any SOAs. However, the mask becomes more disruptive at shorter SOAs, affecting mainly late time of the stimulus processing. The neuronal activity at early time after stimulus onset was previously reported to involve information processing of basic stimulus features (e.g. orientation, motion; Roelfsema et al. 2007). In contrast, neural responses at later time have been reported to involve also feed-back influences and carry information on a figure from background segregation, which is a crucial step for the texture discrimination of the horizontal and vertical stimuli.

### The mask interferes with the figure-ground modulation in V1

The segregation of the target elements from the background in each texture stimulus is a key step towards the discrimination between the horizontal and vertical stimuli. Thus, it is reasonable to assume that the visual system needs to segregate each target from its background, a process that is known as Figure-Ground (FG) segregation. The neural signature of this process in V1 was previously reported in monkeys (Lamme, 1995; Li et al. 2006; Gilad et al. 2013, 2017; Self et al. 2013; Chen et al. 2014; Poort et al. 2016) and can be quantified by the neural response difference between the figure ROI and the background ROI. This response difference was termed as Figure-ground modulation (FGm; Gilad et al. 2013) and to demonstrate its existence in the TDT task we analyzed the V1 response where the animal needed to report the target orientation for trials with stimulus alone but without mask appearance (see Methods). Figure 4A shows a sequence of VSD maps from an example session, for a horizontal stimulus condition. The contour depicts the vertical-only (gray) and horizontal-only (black) ROIs. Following stimulus onset, the VSD maps showed a wide-spread increased neural activation in V1 (map = 100 ms). Interestingly, ∼200 ms after stimulus onset (stimulus duration was 40 ms as in the BM paradigm) the VSD maps showed increased population activity in the horizontal-only ROI, i.e the ’figure’ region for the horizontal stimulus, while the population response in the surrounding background regions, was lower. Specifically, the neural response in the vertical-only ROI, which is part of the background region for the vertical stimulus, was lower. Figure 4B shows the VSD TC of response in the ’figure’ (the horizontal-only ROI) vs. background ROI (the vertical-only ROI) for the data in Fig. 4A. The VSD signal shows an increased activity in the target relative to the background (the response difference between the figure and background is: 1.28×10^-4^±2.51×10^-5^ at early times (80-100 ms; p < 0.01 for difference between figure and background) and the response difference is even larger at later time 5.7×10^-4^±4.3×10^-5^ (180-200 ms; p << 0.001 for difference between figure and background). These results, together with the spatial pattern of activation shown in the VSD maps, reveal the neuronal signature of FGm in V1.

**Figure 4:**
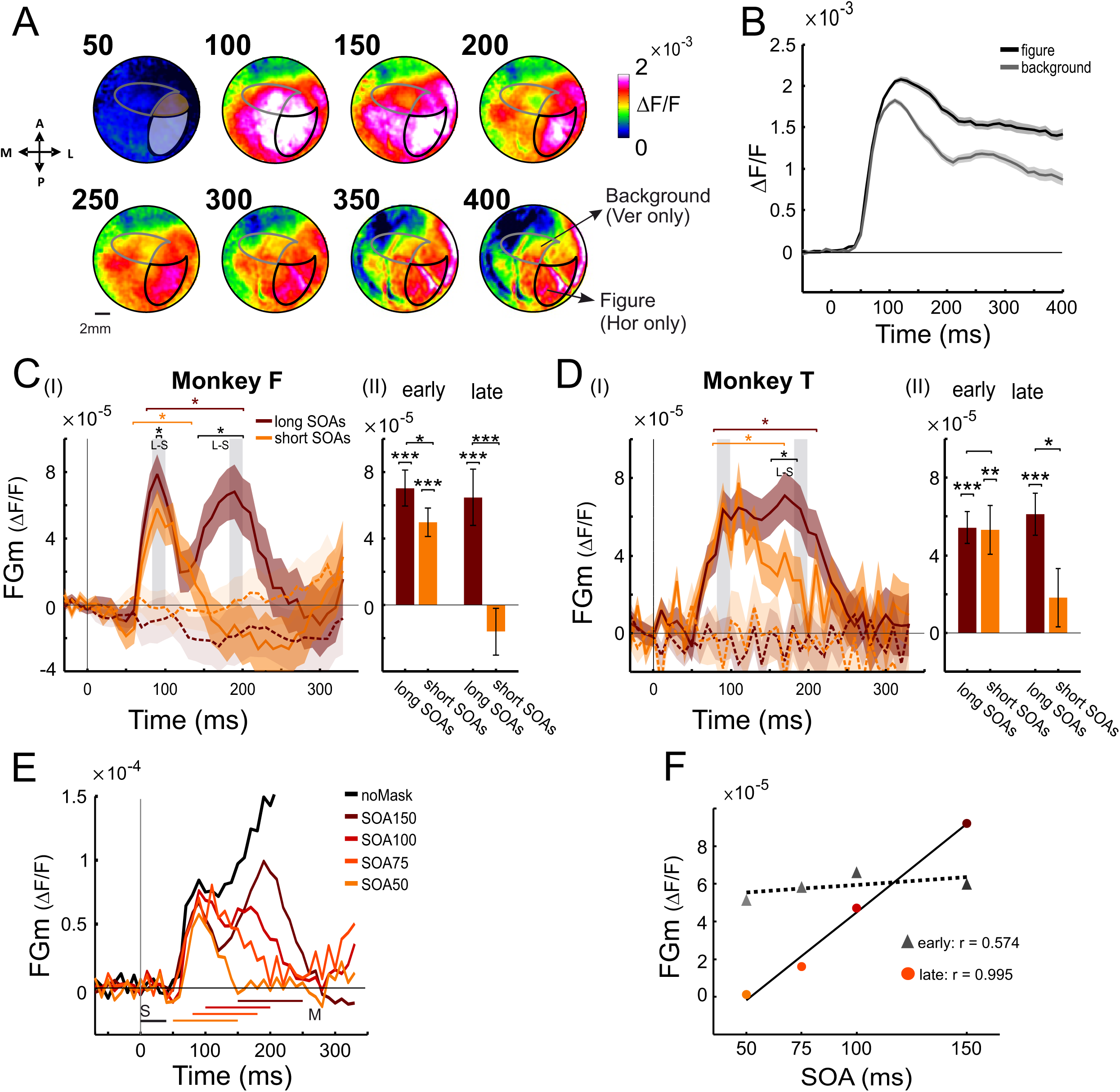
The figure-ground modulation deviates between short and long SOAs at late time. A. Time evolvement of VSD maps for the no-mask condition, stimulus alone (a horizontal stimulus; no mask), an example session (n=29 trials). The contours depict the vertical-only (gray) and horizontal-only (black) ROIs. The late maps show clear population enhancement in the target ROI (i.e. the ’figure’, which is the horizontal-only ROI) relative to the background ROI. B. The VSD TC of response in the target ROI (figure, black curve) and in the background ROI (gray curve) for the same session in A. Note that in the vertical stimulus condition, the black contour represents the figure and gray represents the background, and vice versa in the horizontal condition. C. (i) The time course of FGm from stimulus onset (t=0; see Methods) for long SOAs (150&100 pooled together; n = 115 pairs) and short SOA (75 and 50 ms pooled together; n = 115 pairs), monkey F. Dashed line denotes the shuffled data. (ii) The mean FGm in early (80-100 ms) and late (180-200 ms) times, depicted by gray shaded rectangles. D. Same as C but for monkey T (n = 211 in the long SOAs and 131 in the short SOAs). E. The FGm for the different SOAs, pooled for both monkeys. F. The FGm as function of SOA shows a larger slope (colored circles; r=0.995 p = 0.005) in the second peak as opposed to the first peak (gray triangles; r=0.574; p = 0.43).

Next, we performed a grand analysis (i.e. analysis for all trials) of the FGm in the BM paradigm. In the vertical stimulus trials, we defined the vertical-only ROI as the ‘figure’ ROI and the horizontal-only ROI was defined as the ‘background’ ROI. The opposite was used for the horizontal stimulus trials, where the horizontal-only ROI was defined as the figure and the vertical-only ROI was defined and the background ROI (see Methods; this enabled us to use exactly the same ROIs for figure and background in any of the stimulus). To compute the FGm as function of time, we randomly paired between horizontal and vertical trials, and calculated the population activity difference between the response in the figure ROI and the background ROI respectively. Results were averaged across all trial pairs (see Methods). Next, we pooled together all SOA150 & 100 trials and termed them as ‘long SOAs’ and pooled together all SOA75 & 50 trials, and termed them as ‘short SOAs’. Figure 4Ci shows the TC of the FGm for Monkey F. The FGm shows an early peak around 80-100 ms after stimulus onset, reaching a level of 7.03×10^-5^ ± 1.08×10^-5^ for the long SOAs (p < 0.001, for difference from zero and p << 0.001 for difference between real data vs. label-shuffled data) and 4.98×10^-5^ ± 8.31×10^-6^ for the short SOAs (p=0.025 for difference from long SOA; p < 0.001 for difference from zero; p < 0.001 for difference from label shuffled data). A second peak appears in the long SOAs at around 180-200 ms, reaching an amplitude of 6.45×10^-5^ ± 1.53×10^-5^ which is much larger than -1.67×10^-5^ ± 1.41×10^-5^ for the short SOAs (Fig. 4Cii, p < 0.001 for comparison between long and short SOAs). In monkey T, the FGm as function of time does not show two distinct peaks (monkey B was presented with a lower luminance stimulus; see Methods) but it shows a similar deviation of the long from short SOAs, at late times. In the long SOAs ∼180-200 ms, the FGm reached a level of 6.11×10^-5^ ± 1.18×10^-5^, as opposed to 1.82×10^-5^ ± 1.58×10^-5^ at the short SOAs (Fig. 4Dii, p < 0.01). At early times, the population response in the long SOA reached a peak of 5.43×10^-5^ ± 8.31×10^-6^ between 80-100 ms, and the population response in the short SOAs reached a peak of 5.31×10^-5^ ± 1.28×10^-5^ (Fig. 4Dii, no significant difference between short and long SOAs).

The results of the FGm analysis suggest that the mask interferes with the late FGm processing by significantly reducing it, which might translate to the suppression of targets’ visibility and the ability to discriminate one from each other. The decreased FGm in the shorter SOAs also fits with the monkeys’ reduced behavioral performance when comparing long to short SOAs (an average decrease of ∼12%). Figure 4E shows the FGm response pooled for the two animals (mean across the two animals), separated to the different SOAs. The first peak of the FGm is similar in all SOA, and therefore has very low correlation with the SOA duration and animals’ behavioral performance. At later time from ∼150 to 250 ms, the curves of the FGm in the different SOAs are clearly deviating from each other, and seems to arrange from high to low FGm according to the SOA duration. It is also noticeable that the FGm curve for each individual SOA, shows a decline towards baseline, after the mask onset (plus ∼50 ms delay due to the delay of propagation from the retina to V1). Next, we used a linear fit for the FGm at the early and late peak times, as function of the SOA (Fig. 4F). Interestingly, the late FGm was highly correlated with the SOA duration (Pearson correlation coefficient, r=0.995, p=0.005), while the early FGm was more weakly correlated with the SOA and also non-significant (r= 0.574, p=0.426). In addition, the late FGm had a significantly larger slope when compared to the early FGm (p=0.0012; slopes are: 8.13×10^-8^ and 9.36×10^-7^ for early and late FGm respectively; see Methods).

While the FGm analysis suggested that the mask interferes with the figure-ground segregation processing in V1, it was computed based on the responses in the targets ROIs and also averaged across multiple trials. To examine the contribution of single pixel responses to the stimuli discrimination and investigate the robustness at the single trial level we applied an SVM classifier on the single trials.

### SVM trained on VSD maps successfully classifies trials with a horizontal or vertical target

To verify the applicability of an SVM classifier to the VSD data, we first implemented it on the simple task of discrimination between fixation trials where only a horizontal target or a vertical target were presented to the monkey (see Methods). Figure 5A shows the VSD maps from V1, for an example session in monkey F: Fig. 5Ai shows the population response map evoked by a vertical target and Fig. 5Aii shows the VSD map evoked by the horizontal target (see target stimuli in Fig. 2Ai, 2Bi). The VSD maps were averaged at the time of peak response (70-110 ms after the target onset). Our assumption was that in order to discriminate between the horizontal and vertical targets, the SVM model will converge to opposite weights (i.e. negative and positive weights) for pixels in the non-overlapping targets ROIs.

**Figure 5:**
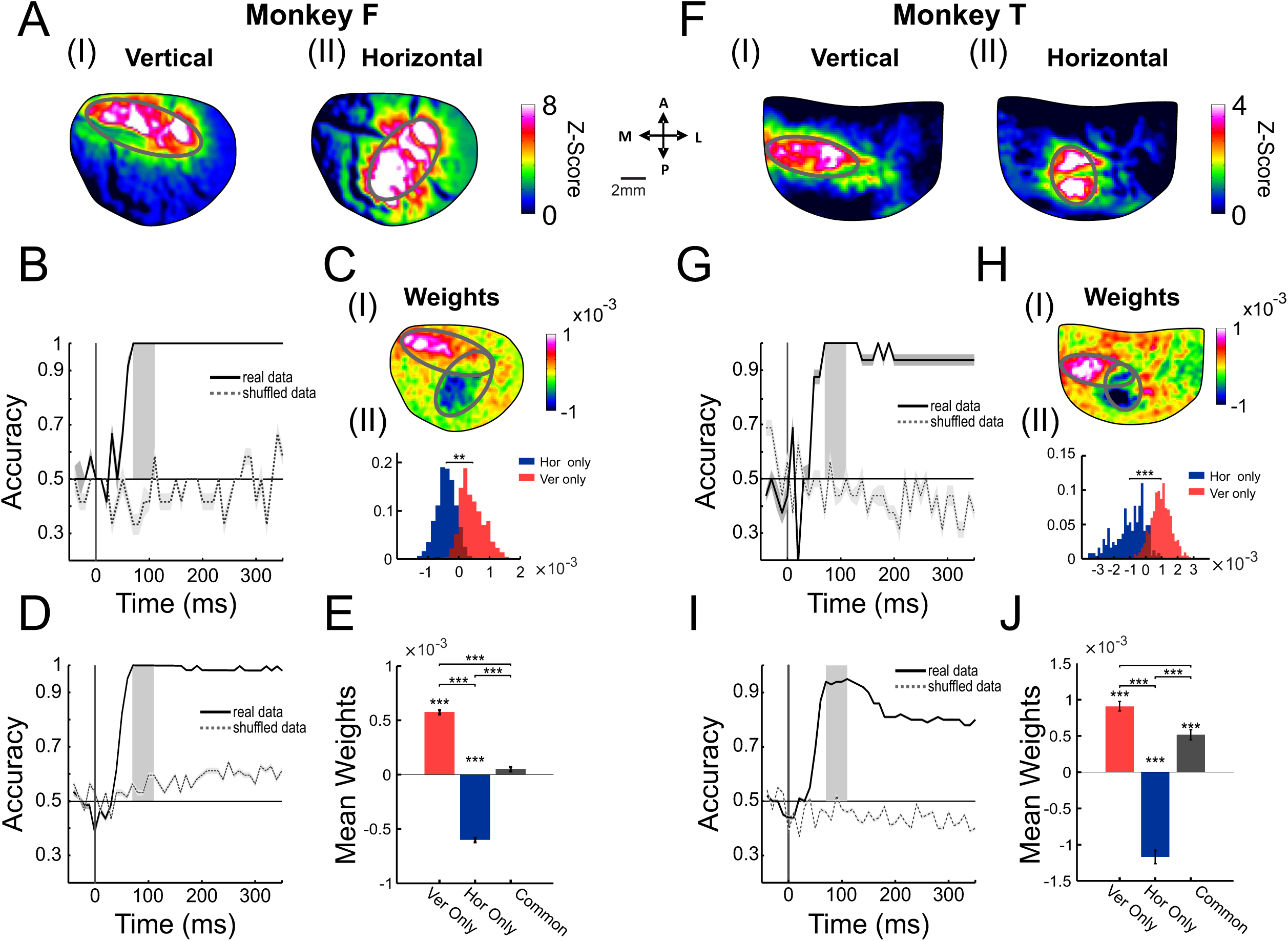
A support vector machine (SVM) classifier on the targets only data: model accuracy and map of weights. Support Vector Machine (SVM) classifier was trained to discriminate between single trials of horizontal and vertical targets (see Methods). **A.** The VSD maps (mean at peak response, 70-110 after stimulus onset) evoked by the vertical (left map; n=6 trials) and horizontal (right map; n=8 trials) targets, an example session for monkey F. **B.** The accuracy of the classifier as function of time (continuous line) and for the shuffled data (see Methods; dashed line) for the data in A. The vertical gray rectangle depicts the time window used for model training (50-100 ms after stimulus onset). Shaded areas around the curves are SEM across all trials. **C (i)** The weights map of the classifier, computed as mean across all model (time window: 50-100 ms after stimulus onset). Warm colors represent positive weights and cold colors represent negative weights. **(ii)** Histogram of the weights in the non-overlapping targets ROIs. **D.** Time course accuracy of the grand analysis model, trained with all trials (n=62). **E.** Mean weights in vertical-only, horizontal-only and common ROI. Error bars are SEM across all pixels. **F-J.** Same as A-E, for monkey T (n=16 trials in condition of the example session; n=100 total amount of trials in the grand analysis).

Next, an SVM classifier was trained with a set of VSD maps (70-110 ms after target onset), from the same example session in 5A. Here each frame in a single trial was used as a single example for the classifier (see Methods). Figure 5B shows the accuracy of the SVM model aligned on stimulus onset (gray rectangle indicates the time window used for training the model). The model accuracy at baseline, before the stimulus evoked response in V1, is near chance level: 51.7±6.01% as expected (mean ± sem; there is no-significant difference between real and shuffled data). However, following the target onset, the model accuracy reached to 100% (within the training window; p < 0.01, rank-sum test between real and shuffled data). Figure 5Ci shows the weight map generated by computing the mean weights across all models (LOOCV, see Methods), where warm colors denote pixels with positive weights and cold colors denote pixels with negative weights. As expected, the largest absolute weights of the model appear at the non-overlapping target ROIs, while in the common ROI of the two targets the weights values are closer to zero. This is demonstrated in the histograms in Fig. 5Cii, showing the weights distribution in the non-overlapping ROIs (Horizontal-only ROI: -4.28×10^-04^±4.31×10^-06^, Vertical-only ROI: 4.59×10^-04^±4.84×10^-05^ (mean ± std), p < 0.001, for the difference between the ROIs; rank-sum test). Figure 5D shows the SVM grand analysis across all target only trials (n= 62, all sessions of monkey F): the model accuracy shows near chance level (p value is n.s. between real vs. shuffled data, rank-sum) before stimulus onset and high accuracy ∼100% after stimulus onset (p < 0.001). Figure 5E shows the grand analysis of the mean weights in the non-overlapping ROIs and in the common ROI. In the horizontal-only ROI the mean weight value is -6.05×10^-04^±1.95×10^-05^ (mean ± SEM) in the vertical-only ROI the mean weight value is 5.73×10^-04^±2.13×10^-05^, and in the common ROI, the mean value is 4.75×10^-05^±2.13×10^-05^. These values were found to be significantly different from each other (p <0.001; rank-sum test).

Figure 5F-J is as Fig. 5A-F but for monkey T. SVM reaches 100% accuracy in the example session (Fig. 5G), and 93.9±2.2% accuracy in the grand analysis model (Fig. 5I). The mean weights value in Fig. 5J in the horizontal-only ROI was -1.16×10^-03^±9.47×10^-05^, in the vertical-only ROI the mean value was 9.10×10^-04^± 6.75×10^-05^, and in the common ROI, the mean value 5.16×10^-04^±7.00×10^-05^ (p < 0.001). These results demonstrate that SVM model is proficient to distinguish between trials evoked by the horizontal and vertical targets, at high accuracy, and the temporal and spatial properties of the model align with the expected results. Interestingly, the SVM weight maps of the target alone trials also suggest the most informative pixels for solving the BM paradigm, because the horizontal and vertical stimuli in the BM task differ only in the non-overlap targets elements that are corresponding to the vertical-only and horizontal-only ROIs.

### SVM of the backwards masking data shows higher accuracy for long SOAs and higher accuracy for correct vs. error trials

Our next step was to apply the SVM model to the VSD data from the BM paradigm, where stimuli are comprised of oriented target elements embedded in a textured background, which is followed by a mask stimulus. A sliding window of 50 ms was used to train and test data (grand analysis across all trials; see Methods). This approach allowed us to generate a time-dependent curve of classification accuracy as function of time from stimulus onset. Figure 6Ai shows the TC of accuracy for the SVM model of long SOAs (SOA 150 & 100) and short SOAs (SOA 75 an SOA 50) in monkey F. Two peaks of accuracy are apparent in both SOAs. The first peak appears during the initial response of V1 activation (∼80-100 ms), which previously was suggested to encode basic stimulus features. Interestingly, a second accuracy peak appears approximately at the same time in which we observed the late phase of the FGm (∼180-200 ms in long SOAs). Figure 6Aii shows that there is a no-significant difference in accuracy between the long and short SOAs during the first peak (0.89 ± 0.01 for long SOAs and 0.88 ± 0.011, no significant difference between the short and long SOAs; p<0.001 for real and shuffled data, rank-sum test). In the late peak, the accuracy for the long SOA is significantly higher than for the long SOA: 0.88 ± 0.015 for long SOAs as appose to 0.79 ± 0.017 for short SOAs; p<0.001 for difference between real data and shuffled data and between long and short SOAs, rank-sum test). Figure 6C shows the same analysis for monkey T with similar results: there is a significant accuracy difference between the long and short SOAs at the late time, although the accuracy values are smaller (see Methods; the stimulus luminance in monkey T was smaller). We hypothesize that the decrease in accuracy for the short SOA at late times is caused by the mask, which may interfere with the stimulus processing and with the FGm.

**Figure 6:**
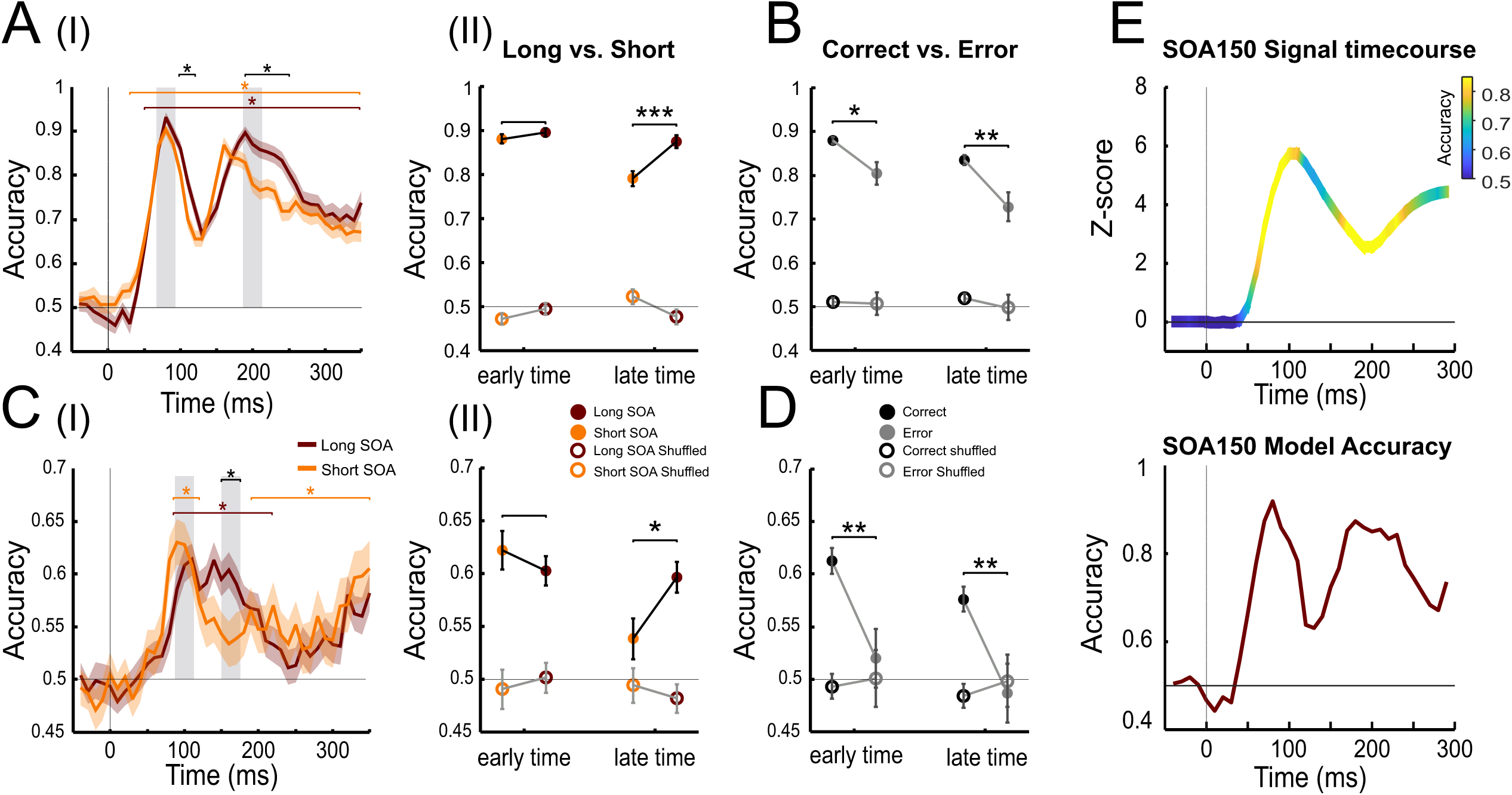
A**n SVM model on the BM trials shows a separation between the long and short SOAs at late times.** An SVM model was trained to discriminate between vertical and horizontal trials using a sliding window of 50 ms (see Methods). **A(i).** Time-course of the model accuracy. Each data-point depicts the mean accuracy across trials (n=234 & 256 trials, for long & short SOA respectively). Dashed lines represent the same analysis except labels were shuffled prior to model training. **(ii).** The mean accuracy at first peak for long and short SOAs (filled circles) and for the shuffled data (empty circles). **B.** The accuracy for error trials is lower as compared with correct trials (filled circles). Error trials were defined as trials in which the animal was presented with a horizontal stimulus and reported on the vertical and vice versa (see Methods; n=72 error trials from all SOAs). Shuffled data appear in filled symbols. **C-D:** Same as in A-B but for monkey T (n=438 & 278 trials, for long & short SOA respectively; n=125 error trials from all SOAs). **E.** Top: Time-course of the VSD signal in V1, for SOA 150. The color coding depicts the model’s accuracy for each time point. Bottom: The model accuracy as function of time for SOA 150 (n=84 trials).

Using the SVM models that were applied for correct trials only, we now tested the model’s accuracy on error trials (see Methods). Error trials were defined as trials in which the animal was presented with a horizontal stimulus, and reported on a vertical stimulus and vice versa. Figure 6B and 6D show that the model accuracy was significantly lower in error trials at both time points, by 8-11% in monkey F (p < 0.05 to p < 0.01) and by 9% in monkey T (p < 0.01). Finally, Fig. 6E top shows the TC of the VSD signal (mean across trials) in V1 as function of time. Superimposed on the TC we added in color coding, the SVM model’s accuracy (see Fig. 6E bottom for the TC of the model accuracy). It is interesting to note that the peak accuracy of the model’s performance does not coincide with the peak response (i.e. highest activity) of the VSD signal. In particular, while the first peak of accuracy is aligned with the rising phase of the VSD signal, the late phase of accuracy is aligned with decreased VSD response after stimulus onset. This result suggests that high level of neural activity does not mean necessarily high accuracy in the model.

### The model weights maps reveal distinct spatial organization for early and late times

To gain insights into how the model solves the BM task, we examined the weights assigned to each pixel in V1, i.e. the weights map (gran analysis across all trials; see Methods). Figure 7A shows the weight maps of the model trained on SOA 150 and SOA 50 trials at early and late times, which are corresponding to the SVM accuracy peaks. The early maps reveal similar patterns for both long and short SOAs. A mixture of positive and negative weight patches is observed across V1 area and also with the targets ROI. These patches are relatively small and may relate to individual elements comprising the texture stimulus or their interactions. Importantly, in SOA150, at the late time, the vertical ROI shows mainly positive weights and the horizontal ROI shows mainly negative weights, indicating a clear spatial arrangement that reminds the SVM weight maps for the target alone stimuli (Fig. 5C). In contrast, the model weight map for SOA 50 shows a less clear pattern. Figure 7B shows the distribution histograms of the pixels weight in the horizontal-only and vertical-only ROIs. The weight difference between the two ROIs is largest for SOA 150, at late time: -1.43×10^-03^±2.45×10^-03^ in the horizontal ROI vs. 1.38×10^-03^±1.98×10^-03^ in the vertical ROI (mean+std), a difference of 2.81×10^-03^ (p < 0.001 for comparison between the two distribution, rank-sum test). Moreover, the area under the curves (AUC; see Methods) shows a value of 0.81 (Fig. 7B, top right, inset). In contrast, at early time of the same SOA, the difference is much smaller: -1.12×10^-03^±4.06×10^-03^ vs. 1.08×10^-04^±3.66×10^-03^, a difference of 1.23×10^-03^ (p<0.001 for the two weight distributions; rank-sum test). In this case the AUC is 0.58, almost at chance level (Fig. 7B, top left, inset). Next, we examined the weight difference for the SOA 50: the weight distribution at early and late time showed similar small weight differences (a difference of -4.62×10^-04^ at early times and - 5.23×10^-04^ at late times, AUC value of 0.476 and 0.433 respectively). Figure 7Ci shows the weight difference, pooled for the two monkeys as function of time. While at early time (80-100 ms) there seems to be a small separation between the SOAs, in the late time (180-200 ms) there is a clear separation, revealing that longer SOAs show a larger difference between the weights in horizontal-only and vertical-only targets. Figure 7Cii shows a linear fit for the weight difference of early and late times as function of SOA. While both first and late peak show high correlation values, only that of the late peak is statistically significant (r=0.99, p=0.009). Moreover, there was a significantly larger slope for the weight difference at late time as compared with the early time (p=0.015; slopes are: 1.22×10^-5^ and 3.33×10^-5^ for early and late weight difference). Figure 7Ciii shows the AUC analysis for the weights’ distributions in the early and late time. To further examine whether the weight maps is similar to the weight maps for the target alone trials (see Fig. 5C, 5H) we computed the correlation between the weight maps of the different SOAs and the weight maps obtained from the target alone trials. The linear regression of the correlation as function of time appears in Fig. 7Di and shows high similarity to the results shown in Fig. 7Ci. The correlation analysis shows a larger separation between the SOAs at late time with the highest correlation values for SOA 150 and lowest for SOA50. Moreover, Fig. 7Dii shows that the correlation at late times was strongly dependent on the SOA (r=1, p=0). In addition, there was a significantly larger slope for the correlation at the late time as compared with the early time (p=0.00078; slopes are: 6.88×10^-4^ and 3.31×10^-3^ for early and late correlation). We suggest that the spatial pattern at the late time reflects a top-down feed-back influence into V1, that encodes the late FGm, representing the target and their segregation from the background region.

**Figure 7:**
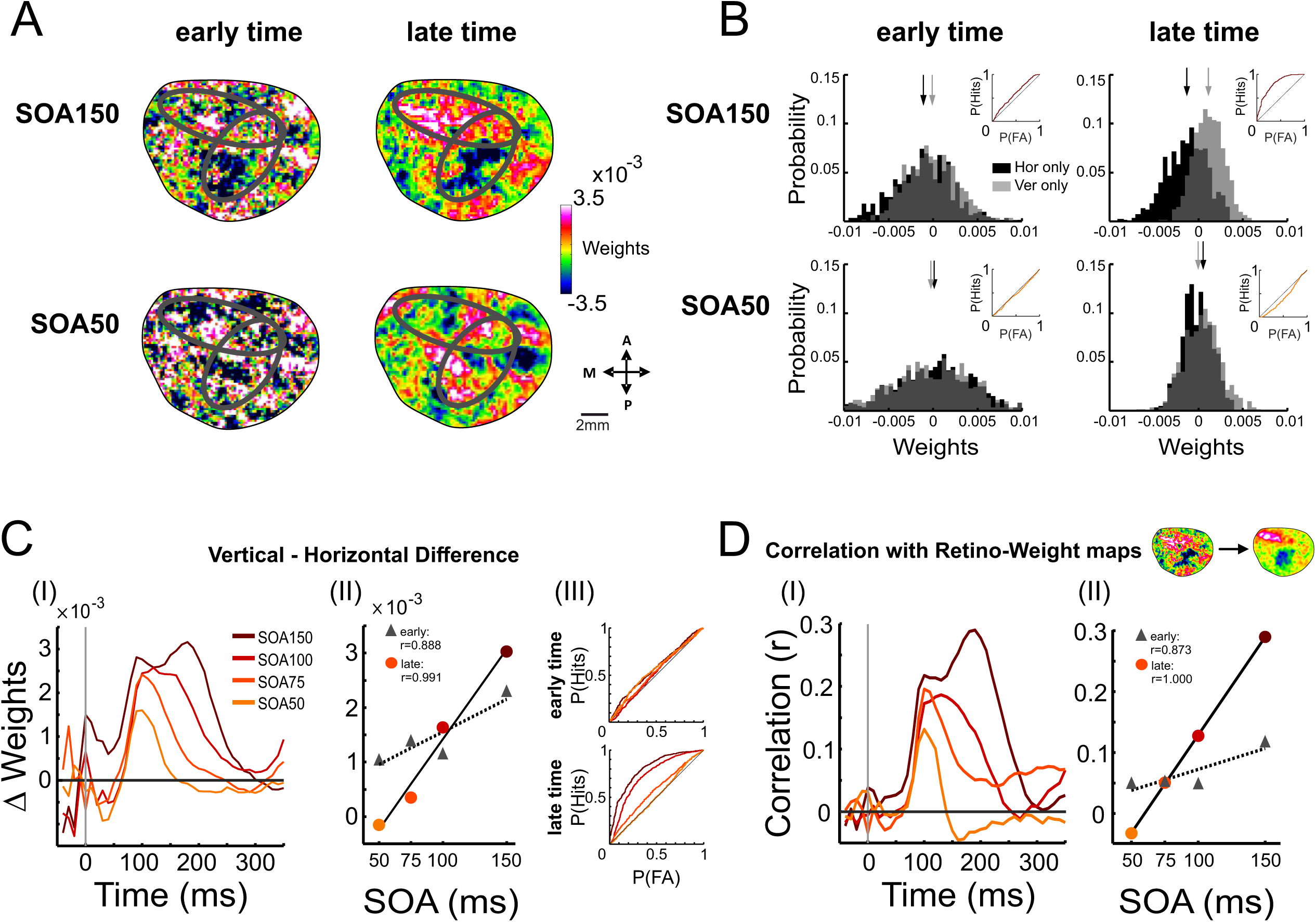
T**h**e **weight maps of the SVM classifier in long and short SOAs A.** Weight maps from the grand analysis model, for early and late time-window (time window: 70 ms & 180 ms after stimulus onset, respectively), for SOA 50 & 150. **B.** The weight histograms in the Ver only and Hor only targets ROIs (n = 604 & 617 pixels in Hor only and Ver only targets, respectively). Insets (top-right in each panel) is the Receiver Operating Characteristic curve corresponding to the histograms below. The area under the curve (AUC) values are: 0.585 and 0.81 for early and late time in SOA 150 and 0.48 and 0.43 for early and late time in SOA 50. **C (i)**: The weights difference between the vertical-only and horizontal-only ROIs, as function of time, for the different SOAs. **(ii).** The weights difference as function of SOA in the first (r=0.88, p=0.112) and second peak (r=0.99, p=0.009). **(iii)** The ROC curves for the weight distributions in the horizontal-only and vertical-only ROIs, for the different SOAs, in the early and late time. **D.** Left: The correlation between the target alone-weight maps (Fig. 3) and the weight maps from the BM, as function of time. Right: The correlation between the weights maps as function of the SOA. First peak: r=0.873 p=0.127, second peak: r=1, p=0.

### Classification analysis using a neural network model supports the SVM results

To examine whether the SVM results on the VSD data can be replicated using a different approach, we used a convolutional neural network (CNN) model (see Supplementary material). Unlike SVM, which treats each pixel as an independent feature, CNNs are better suited for image classification as they capture spatial patterns and spatial correlation between the image pixels. We used the same VSD dataset and time windows as used for the SVM, and first tested the applicability of the CNN on the VSD data using the target alone data. The accuracy of the CNN model on the target alone data showed high values following stimulus onset (99.7% and 83% for monkey F and T respectively; Supplementary material and Fig. S2A, S2B). Analysis of feature importance (SHAP analysis, see Methods in supplementary material) confirmed the strong contribution of vertical ROI pixels and horizontal ROI pixels for the classification task in both animals. For monkey F, the mean SHAP value is: -1.71 ± 0.04 (mean ± SEM) and 1.42 ± 0.033 in the horizontal-only ROI and vertical-only ROI, accordingly. In the common ROI the SHAP value is 0.04 ± 0.035. These values are significantly different between them (p << 0.01, rank-sum test). A similar pattern is observed in monkey T, with mean SHAP values of -1.52 ± 0.07 in the horizontal ROI, 1.67 ± 0.06 in the vertical ROI, and -0.137 ± 0.09 in the shared area (p << 0.01, Wilcoxon rank-sum test). In the BM data, the TC of model accuracy for horizontal and vertical trials showed temporal pattern that was similar to the SVM model: an early peak followed by a late peak in time (Fig. S2C in supplementary material). Using the CNN model, we confirmed a deviation between short SOA and long SOAs at late time, with higher accuracy for the long SOAs. The SHAP analysis revealed similar spatial patterns to those of the SVM (Fig. S2D in Supplementary material).

## Discussion

Using VSDI we investigated the V1 population responses in pattern BM. The neural activation revealed a FGm that was lower for short SOAs in comparison with the long SOAs. An SVM classification revealed differences in the temporal dynamics of the model accuracy in relation to stimulus visibility and spatial reorganization of the informative pattern of activation in long vs. short SOAs. Our results suggest that the mask interferes with the late neuronal activity in V1, that is corresponding to the FGm in V1and can lead to reduced stimulus visibility.

The monkeys’ behavioral performance at shorter SOAs was reduced (Fig. 1), which is in accordance with previous studies in human (Karni and Sagi, 1991; Scoups and Orban, 1996; Censor et al., 2009) and in monkeys (Kovacs et al. 1995; Thompson & Schall 1999; Lamme et al. 2002; Rolls 2004; Op de Beeck et al. 2007), in rodents the behavioral effects in BM are under debate (Dell et al. 2019). In addition, the monkeys’ RT increased as the SOA shortened, suggesting that a weaker stimulus visibility requires extended processing times for report (and may reflect also a lower confidence of the taken decision). These results are consistent with a study in humans by Bacon-Mace et al. (2005), who showed that RTs were significantly slower in the short SOAs in comparison with the long ones (see also Schubert et al. 2013; Bacon-Macé et al. 2007). A few other studies have investigated the relationship between stimulus visibility and RTs in monkeys, using tasks such as contrast detection (Palmer et al., 2007) or motion detection (Cook & Maunsell, 2002). These studies reported on longer RTs under conditions of reduced stimulus visibility. These observations are aligned with our findings, where shorter SOAs were associated with longer RTs and reduced visibility (i.e. poorer performance rates).

We found that the V1 responses to the mask alone and stimulus alone showed an increasing overlap in time as the SOA became shorter, while the early phase of the response to the stimulus was unaffected (Figure 3C). These results are in accordance with a previous ERP study in humans (Censor et al. 2009) and suggest that while the feedforward transfer of stimulus information to V1 was preserved at all SOAs, the mask overlap and thus interference with the stimulus processing was larger in short SOAs. By precisely mapping the different stimulus parts, i.e. the targets (the ’figures’) and background regions onto the V1 surface, we computed the FGm in the BM paradigm and studied the effect of the mask on the FGm for the different SOAs. Interestingly, the TCs of the FGm for short and long SOAs showed a deviation at late time (∼ 200 ms after stimulus onset), with higher FGm values for long SOAs. This observation was consistent with the no mask condition, that revealed the spatial signature of FGm (Fig. 4A; increased activity in the target ROI and decreased activity in the background ROI) and showed an elevated FGm from early to late time.

The emergence of FGm at late time was previously reported (Lamme, 1995; Li et al. 2006; Gilad et al. 2013, 2017 using VSDI; Self et al. 2013; Chen et al. 2014; Poort et al. 2016) along with additional studies that described an early time component of the FGm. The latter appears during the rising phase of the stimulus evoked activity and relates to information processing of basic stimulus features (e.g. orientation, Roelfsema et al. 2007). Our results show that the mask did not interfere with the early phase of FGm (up to ∼ 100 ms after stimulus onset; Fig. 3C, 4E, 4F), however it disrupted the late FGm (Fig. 4E, 4F). Moreover, we found a higher correlation and larger slope between the FGm and the SOA duration for the late FGm as compared with the early FGm. This is in accordance with Lamme et al. (2002), who recorded the spiking activity of V1 neurons in monkeys and showed that the FGm signals are selectively and strongly suppressed at SOAs that induced invisibility of the stimulus. Several studies have proposed that FGm depends on feedback from extra-striate areas (Lamme, Supèr and Spekreijse 1998; Li et al. 2006; Gilad et al. 2013, 2015; Chen et al. 2014; Poort et al. 2016). Therefore, one can suggest that the masking interrupts the recurrent interactions between V1 and higher visual areas.

To investigate the most informative pixels in V1 for the BM task, at the single trial level, we applied an SVM classifier for each time-point along the VSD TC. We used the SVM decoding accuracy to quantify the amount of task related information that existed in V1 as function of time. The accuracy for the targets alone stimuli increased at the rising phase of the population response to the stimulus and persisted in V1 long after the stimuli was turned off, suggesting that the residual activity in V1 continue to hold information about the stimulus features. The weight maps showed, as expected, that the neural activity in the non-overlapping target ROIs carry the information to distinguish between the two stimuli. This analysis showed that SVM is proficient in discriminating single trials of horizontal and vertical targets and suggested also the spatial solution for solving the BM paradigm. In the BM task, the decoding level at early time were similar across SOAs, however, the accuracy at the late time revealed difference between long and short SOAs that fits with the behavioral report on stimulus visibility (Fig. 6A, 6C). We suggest that the decoding accuracy at early time, reflects the feed-forward information and V1 local processing of the elements’ orientation (Lamme et al. 2002; Gilad et al. 2013; Roelfsema et al. 2007). The late time of accuracy seems to reflect feedback influences from higher brain regions that drives the late FGm and was shown to be linked with behavioral report (Lamme, Supèr and Spekreijse 1998; Lamme et al. 2002; Li et al. 2006; Gilad et al. 2013, 2015; Chen et al. 2014; Poort et al. 2016; Gilad et al. 2013, 2015). Additional analysis showed a lower level of accuracy for the error trials relative to the correct trials, in both the early and late time phases. This result suggests that the error trials may emerge due to degraded neural activity either at the early or late time of information processing in V1. The first one may emerge from impaired feedforward transfer of stimulus information and its processing and the late one may result from disruptions in feedback influences.

Interestingly, an analysis of the SVM weight-maps revealed for long SOAs (where stimulus visibility is better) a spatial re-arrangement of the weights as function of time from stimulus onset. In the early time of the neural response, the weight map showed multiple distributed clusters all over V1, whereas at the late time the weights tended to accumulate in the target ROIs, with opposite sign in the two target ROIs. The ROC analysis in Fig. 7B confirmed that the weights distributions showed a higher separation at the late time of the long SOA. Figure 7C showed that the weight difference as function of time was larger at late times and also the correlation with the SOAs duration was higher (along with a larger slope for the linear regression fit) the for the late time as compared with the early time. In addition, there was a higher correlation between the weight maps at late time to the weight maps obtained for the target alone trials. Interestingly, the weight maps of the target alone trials also suggested the pixels that are most informative for solving the BM paradigm: because the horizontal and vertical stimuli differ only in the non-overlap target elements reflecting the vertical-only and horizontal-only ROIs. Therefore, the SVM model uncovered the spatial rearrangement of the task-related informative activity in V1, and this spatial rearrangement matched the non-overlapped target ROIs. This rearrangement of the weights did not occur for short SOA, which suggest that the mask indeed disrupts the top-down influences.

In summary, our results support the view that feedforward processing alone is not sufficient for stimulus percept and offer a better support to the theory of disrupted cortical feedback in BM. While decoding levels during the early phase were high in all SOAs, the behavioral performance of the animal was substantially lower in short SOA, despite all information being represented in V1. We found that the animal behavioral performance was better correlated with the late phase of FGm and SVM at late time (accuracy and weight maps), which we suggest to be driven by top-down influences and is susceptible to mask interruption.

## Methods

### Behavioral task and visual stimuli

Two adult monkeys (Macaca fascicularis; F and T, 9 and 12 Kg) were trained on a texture discrimination task (TDT) with pattern BM as previously used in human subjects (Karni and Sagi, 1991). The stimuli were computer generated textures, made of short line segments (Fig. 1Ai). Visual stimuli were presented on a 21-inch Mitsubishi monitor at 85 Hz, placed 1 meter from the monkey’s eyes and occupied 10°×10° (line segment length: 0.5°, width: 0.1°, mean distance between line elements’ center: 0.75°). A horizontal or a vertical stimulus refers to a horizontal or a vertical target consisted of three horizontally or vertically aligned lines, that were embedded in a texture background (Fig. 1Ai left and middle). The targets were segregated from the background elements by their orientation difference only. Background elements were jittered randomly (±0.1-0.3°). The mask stimulus (same size as the stimulus) was made of random oriented V-shaped elements (Fig. 1Ai right). The luminance of all elements (target, background and mask elements) was 74 cd/m^2^ for monkey F and 23 cd/m^2^ for monkey T. The background luminance for both monkeys was near zero. The luminance levels of the stimuli enabled us to investigate whether the neural modulations are preserved across variable luminance conditions of the stimulus.

A trial started when the animal fixated on a small fixation point. After a random time interval (3000-4000 ms) a horizontal or vertical stimulus appeared on the screen for 40 ms. Then, after a variable period of time, a mask appeared for 100 ms (Fig. 1Aii). The time interval between the stimulus onset to mask onset, is defined as stimulus-to-mask onset asynchrony (SOA). Following the mask offset (the fixation point and mask were turned off together), two small lateral points appeared and the monkey had to report the target orientation (horizontal or vertical) by making a saccade either to the right or the left, within a brief time window (500 ms). A trial was classified as correct, only if the monkey reported the target orientation correctly. The animal was rewarded with a drop of juice or water for each correct trial. The stimulated trials were interleaved with fixation-only trials (blank condition; no stimulus was presented). This condition was used to subtract the heartbeat artifact (see basic VSD analysis) and was used as baseline activity condition.

The animals were also trained on a simple fixation task, where they were required to maintain fixation throughout the entire trial (no report was requested) and were rewarded for holding the fixation. On each recording day, the animals performed a fixation task where a stimulus comprised of the horizontal or vertical target alone was turned on (Fig. 2Ai, Bi). The duration of the target stimuli was identical to the stimulus duration in the BM task. This approach enabled us to identify in V1 the regions evoked by the targets alone and background alone, i.e. to retinotopically map the targets and background regions onto the imaging V1 area (Fig. 2A-2C). In another set of experiments the monkeys performed a fixation task, while they were presented with the stimulus alone image (i.e the texture stimulus: target embedded in the background as shown in Fig. 1Ai; Fig. 3A shows the VSD response to the stimulus alone) or the mask alone stimuli (Fig. 1Ai; Fig 3B shows the VSD response the mask alone). The stimulus duration was as in the BM task and mask onset time and duration was identical to the different SOAs conditions in the BM paradigm. These imaging sessions served as control conditions, to measure the VSD signal evoked by either the stimulus alone or mask alone (Fig. 3) and enabled us to compare the V1 activity in the BM task to the VSD signal in the stimulus and mask alone conditions (Fig. 3E). In another set of experiments, the stimulus alone appeared (for 40ms) without the mask (Fig. 4A, 4B, 4E) and the monkey needed to report which stimulus it perceived, i.e. a vertical or horizontal stimulus, by performing a saccade either to the right or the left. The datasets for all behavioral paradigms are detailed under ’VSDI dataset and analysis’.

### Data Acquisition

Two linked computers controlled the visual stimulation, data acquisition, and the monkey’s behavior (CORTEX software package). The protocol of data acquisition in VSDI has been described in detail elsewhere (Slovin et al., 2002). Single trials were saved on separate data files to enable single trial analysis.

### Eye-position recording

Throughout the experimental paradigm, the monkey’s eye positions were monitored by an infrared eye tracker (Dr. Bouis Device, Kalsruhe, Germany), sampled at 1 KHz and recorded at 250 Hz. Only trials with tight fixation, where the animal maintained within ±1 deg window until the report time (in the TDT with or without the BM) or until end of trial (in the simple fixation paradigms), were used for the analysis.

### Behavioral analysis

To study the behavioral performance in the BM experiments, we computed the performance of the animal as the percent correct trials in each session, i.e. the fraction of correct trials relative to the total number trials (correct and error trials) in the same session. Trials with early break fixation (the monkey did not keep fixation until the mask and fixation point offset) were excluded from analysis. A correct trial was defined as a trial in which the monkey identified correctly the target’s orientation and reported within 500 ms whether the target was vertical or horizontal by a saccade. Error trials were defined as trials with wrong responses, i.e. a horizontal stimulus was presented but the animal reported on a vertical stimulus and vice versa. We computed the percentage of correct trials for each SOA session separately and created a psychometric curve for each monkey (Fig. 1B; Monkey F: n = 8, 6, 8 & 6 sessions, Monkey T: n = 6, 12, 10 & 12 sessions for SOAs 50, 75, 100 & 150 ms respectively). We also computed the reaction time (RT) of the monkeys that is defined as the time interval from fixation point and mask offset to the onset of the reporting saccade. The RT analysis was done at the single trial level for correct trials (Fig. 1C; Monkey F: n = 173, 111, 171, 106 trials for SOAs 50, 75, 100 & 150 ms respectively. Monkey T: n = 124, 354, 358 & 399 correct trials for SOAs 50, 75, 100 & 150 ms respectively). A RT analysis was done at the single trial level also for error trials and compared to the correct trials (Supplementary Fig. S1). This single trial analysis at the behavioral level was done to fir with the grand analysis of the VSD data was done on the pooled dataset of single trials. While it is possible to compute RT at the single trial level, it is not possible when computing percentage of correct trials, which is done at the session level.

### Surgical procedure and VSD imaging

All experimental procedures were carried out according to the NIH guidelines, approved by the Animal Care and Use Guidelines Committee of Bar-Ilan University, and supervised by the Israeli authorities for animal experiments. Antibiotics and analgesics were applied before, during and after surgical procedures and adequate measurements were taken to minimize pain and discomfort. The surgical procedure has been reported in detail elsewhere (Arieli et al., 2002; Slovin et al. 2002). Briefly, a craniotomy was performed under full anesthesia and aseptic conditions, and the dura mater was removed, exposing the visual cortex. Then, a thin and transparent artificial dura made of silicone was implanted over the visual cortex, to enable long-term imaging. The anterior border of the imaged area was typically 3-6 mm anterior to the Lunate sulcus. For the retinotopic position of the imaged V1 see ’Retinotopic mapping of the V1 area and the border between V1 and V2’.

We stained the cortex with voltage-sensitive-dyes: RH-1691 or RH-1838 supplied by Optical Imaging, Israel. VSDI was performed using a MiCAM Ultima system with sampling rate of 10ms/frame (100Hz) and a spatial resolution of 10,000 pixels/frame. Each pixel sums the neuronal activity from an area of 170^2^ µm^2^ that includes a few hundred neurons. The exposed cortex was illuminated using an epi-illumination stage with an appropriate excitation filter (peak transmission of 630 nm, width at half-height of 10nm) and a dichroic mirror (DRLP 650), both from Omega Optical, Brattleboro, VT. A barrier post-filter above the dichroic mirror (RG 665, Schott, Mainz, Germany) collected the fluorescence and rejected stray excitation light.

### Retinotopic mapping of the V1 area and the border between V1 and V2

For the retinotopic mapping of the imaged V1 and the border of V1/V2, we used we used 6 sessions of optical imaging of intrinsic signals (OIIS) in monkey F and 7 sessions OIIS in monkey T. After the animal acquired fixation and maintained fixation for a random time interval (2-3 s), a stimulus appeared over the screen for 2 s. The animal was required to maintain fixation for additional 4 s after the stimulus offset. The stimuli were a 0.5° vertical bar or horizontal bar of moving square gratings. The bar position varied at increasing distances (deg) from the vertical meridian (VM) or from the horizontal meridian (HM), correspondingly. In Monkey T, the center of the imaged chamber was ∼2.25° deg below the horizontal meridian and 1° from the vertical meridian. In Monkey F, the center of the imaged chamber was ∼1.5° below the horizontal meridian and ∼2° from the vertical meridian. To map the border between V1 and V2 we performed OIIS and obtain the orientation domains and ocular dominance maps. Using these functional domains, we could map the border between V1 and V2 areas (Shtoyerman et al. 2000; Slovin et al. 2002).

### VSDI dataset and analysis

VSDI data was obtained from the right hemispheres of two adult monkeys (Macaca fascicularis; F and T, 9 and 12 Kg). A total of 27 and 28 VSDI sessions were analyzed in monkey F and T respectively (11 imaging days). This dataset is a little smaller than the behavioral dataset because it includes removal of VSD trials with high noisy (Gilad et al. 2013) and matching trial numbers between vertical and horizontal stimuli for SVM and FGm analysis. The dataset for VSDI retinotopic mapping of the target and background regions in V1 (Fig. 2) was comprised from 62 trials (4 sessions) in monkey F, and 100 trials (6 sessions) in monkey T. The same dataset was used for training the SVM model of the target alone trials, i.e. the retinotopic data (Fig. 5). The VSDI dataset for BM paradigm was comprised from: Monkey F: 14 sessions that included 158, 98, 150, 84 trials for SOAs 50, 75, 100 & 150 ms respectively. Monkey T: n = 20 sessions that included 70, 208, 190 & 248 correct trials for SOAs 50, 75, 100 & 150 ms respectively (Figs. 3,4,6,7). The dataset for the stimulus alone and mask alone was comprised of 33 trials for stimulus alone (3 sessions) and 29, 26 & 15 trials for the mask alone stimulus (5 sessions), using onset times that were matched to the mask in those in the BM task: SOAs 50, 100 & 150 ms respectively (Fig. 3). For the example session of the TDT paradigm without the mask, we used 29 trials from one session in monkey F (Fig. 4A, 4B). For the grand analysis of the no mask condition in the FGm analysis (Fig. 4E) we used 26 trials (from 3 sessions) for monkey F, and 86 trials (2 sessions) for monkey T. For the error analysis in Fig. 6B (comparison of SVM accuracy in error and correct trials) we used a total (short and long SOAs) of 72 and 125 error trials in monkey F and T respectively.

### Basic VSDI analysis

All data analyses were done using MATLAB software. The basic analysis of the VSDI signal is detailed elsewhere (Slovin, 2002; Ayzenshtat et al., 2010). Briefly, to remove the background fluorescence levels, each pixel was normalized to its baseline fluorescence level (average over few frames before stimulation onset). The heart beat artifact and the photo bleaching effect were removed by subtraction of the mean blank signal (fixation only trials) from stimulated trials. Thus, the imaged signal (Δf/f) reflects the changes in fluorescence relative to the blank trials. The VSD maps in Fig. 2,3,4,5 were low pass filtered (2D Gaussian filter, sigma=1 pixel) for visualization purposes only.

### Defining regions of interests (ROIs) for the targets and background V1

On each recording day, the animals performed a fixation task (see Behavioral tasks and visual stimuli) where a horizontal alone or vertical target alone was turned on (for 40 ms, as in the BM task). This approach enabled us to retinotopically map the targets and background regions onto the imaging V1 area (Fig. 2). To study the population response in the target and background regions and their relations in the various behavioral tasks, we defined ROIs for each region. First, we computed the activation patterns evoked by the target alone conditions (Fig. 2A, 2B) to define separate ROIs for vertical and horizontal targets by fitting an elliptical shape ROI to the activation map of each target (80-100 ms after stimulus onset). Figure 2Aii shows the ROIs for the targets: vertical ROI (ver-ROI) and Figure 2Bii shows the horizontal ROI (hor-ROI). Figure 2Cii shows the background and the common target ROI as well as the non-overlapping parts of the target ROIs: ver-only ROI and hor-only ROI.

### Analysis of the overlap between the stimulus and mask responses and mask suppression

To compute the overlap between the stimulus alone evoked response and the mask alone evoked response in V1, we first averaged the response across V1 for all trials in each condition and session, separately. An example session is shown in Fig. 3A-B. Next, the TC of each single trial was normalized to the peak response of the trial-averaged signal (of the same condition and the session), a normalized example session appears in Fig. 3C. The shaded area (orange-pink) in Fig. 3Cii depicts the overlap between the stimulus alone response and mask alone response. This was computed as the area underneath the joint curves of the stim and mask alone responses, from mask onset until t=230 ms after stimulus onset. Figure 3Ciii depicts the grand analysis of this response overlap, which was computed for all trials in all sessions. To compute the mask suppression (Fig. 3E), we plotted the VSD TC from an example session of each BM SOA with its corresponding mask alone stimulus (i.e., SOA 150 with Mask onset 150). We then identified the two intersections of the mask alone curve with the BM curve (orange-pink area) and calculated the area locked between them by subtracting the area under the BM curve from the area under the mask alone curve between the two intersections points, for all the trials.

#### Figure-ground modulation

The figure-ground modulation (FGm) is defined in equation no. 1:

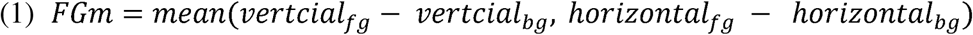

The FGm is calculated as the response difference between the figure and background regions for pairs of horizontal and vertical trials (taken from the same imaging session). In each trial, the ‘figure’ ROI was defined as the region activated by the target elements of that orientation, excluding the ‘common element’ region (i.e. vertical_fg_ and horizontal_fg_). The ‘background’ ROI (vertical_bg_ and horizontal_bg_) was defined as the region which the opposite target activated (excluding the ‘common element’). Thus, for the vertical trial, the vertical_fg_ and the vertical_bg_ are the vertical-only and horizontal-only ROIs and for the horizontal trial the opposite applies, the horizontal_fg_ and the horizontal_bg_ are the horizontal-only and vertical-only ROIs (Fig. 2Cii, Fig. 4A). After this calculation, all FGm pairs from all recording sessions were pooled together according to their SOA-condition (Fig. 4C, D, E). This approach for computing the FGm enabled us to use exactly the same brain regions for computing the FGm in both horizontal and vertical stimuli and thus account for any variation of VSD staining across recording days.

To estimate the expected FGm by chance, we randomly shuffled the trials label within an SOA session, and assigned vertical/horizontal labels such that the ROIs now represent ‘figure’ and ‘background’ regions at random. This control condition showed a near zero value of the FGm (Fig. 4C, D dashed lines). To assess the significance level of the results, we performed the label shuffling 100-fold, and created a distribution of this shuffled FGm. We then performed a Wilcoxon rank-sum Test with the values of the real data vs. the shuffled ones.

### Spatial registration of the VSD maps between different recording days

The cortical images obtained from the different recording days showed slight variations in the exact chamber location and orientation relative to the camera frame. This occurred due to slight changes in camera position relative to the imaging chamber. Accounting for these variations was crucial for effective training of the linear SVM model, where each pixel is a feature and across the recording days a given pixel should represent the same site in V1. To address this, we selected one recording day as a fixed reference and registered all other recording days to it by aligning the blood vessels pattern of the cortical tissue in each session with those of the reference session (in a previous study we showed that the functional columns in V1 remain constant in space relative to the blood vessels, Slovin et al. 2002). In addition, for monkey T, there was a minor shift (0.5°) of the stimulus position on the display between the first 3 recording days and the last 3 days days. We therefore divided the data into two datasets of 3 sessions each, that were separately registered and aligned. The SVM analysis was performed on the VSD registered maps and all results were verified on original data (i.e. before registration).

### An SVM model on the target alone data: accuracy and weights map

For the SVM analysis, we normalized the VSD data relative to baseline noise. This was done by normalizing the activity in each pixel to Z-score, with the mean and standard deviation calculated from baseline frames, i.e. before stimulus onset (220 to 70 ms before stimulus onset, from all trials of a single session). This normalization is a standard approach for SVM analysis.

For training the SVM on the VSD maps of the target alone data, we used a training window of 70-110 ms (i.e. 5 frames) after stimulus onset, as this is the time when the stimulus evoked activity becomes detectable in the VSD maps. We balanced the training data by selecting an equal number of trials from the horizontal and vertical conditions – 12 & 16 trials total for monkeys F and T respectively in the example sessions analysis (Fig. 5B, C, G, H) and 62 & 100 trials total for monkeys F and T respectively in the grand analysis (Fig. 5D-I). The Leave-One-Out Cross-Validation (LOOCV) method was employed, meaning that in each training iteration, one horizontal and one vertical trial were left out for evaluation while the remaining trials were used for training the model. The 5 frames of the training window from each trial were pooled together across all training trials and used to train the SVM. We then tested the model on the two trials left out for each time point and this procedure was repeated for all pairs of trials. Next, the model’s performance was averaged across all pairs of trials to create the accuracy TC shown in Fig. 5. This entire process was conducted separately for monkeys F and T, initially using an example session (Fig. 5B, 5G) and later expanding to the entire dataset (Fig. 5D, 5I). Finally, we averaged the weights of all trained models to produce one single weight map, and color-coded it such that warm colors represent high positive weights and cool colors represent low negative weights (Fig. 5Ci, 5Hi). Using the horizontal and vertical ROIs that were defined previously, we calculated the weight distributions (Fig. 5Cii, 5Hii). The mean weights in the vertical-only, horizontal-only and common ROIs of the grand analysis is shown in Fig. 5E, J. When training the data on the SVM we equalized the number of trials for the horizontal and vertical stimuli.

To assess the statistical validity of the SVM model, we performed label shuffling on the training set before training. We then used the Wilcoxon rank-sum test to compare the distribution of model performances within the training window between the shuffled and unshuffled data, as well as to compare the mean weight values of the ROIs within the weight maps.

### SVM on backward masking data: accuracy and weights map

To investigate the temporal dynamics of the model accuracy with the BM data, we used a sliding window of 5 frames (50 ms) with a stride of 1 frame (10 ms). For each window the SVM model was trained on the VSD maps of the BM data using a LOOCV approach (as described above in ’An SVM model on the target alone data: accuracy and weights map’) and tested using a 5-frame window on one pair of trials: one horizontal and one vertical stimulus. Model accuracy was averaged across these frames and all trials. Each SOA was trained and tested independently, allowing for a comparison of the model’s performance across different SOAs (Fig. 6). When training the data on the SVM we equalized the number of trials for the horizontal and vertical stimuli. To assess the statistical significance of the results, we performed label shuffling of the trials prior to training and used Wilcoxon rank-sum test to compare the real data to the shuffled data. Rank-sum was used to compare models’ accuracy across the different SOAs at the early and late time of accuracy.

Finally, we averaged the weights of all trained models to produce one single weight map that is color-coded (Fig. 7A) as for the SVM on the target alone data. The weight maps were low pass filtered (2D Gaussian filter, sigma = 0.5 pixel) for visualization purposes only. Using the horizontal and vertical ROIs that were defined previously, we calculated the distribution histograms of the weights in the vertical-only and horizontal-only ROIs (Fig. 7B). To measure how much the weights distributions are different between the ROIs, we calculated the ROC curves and the AUC between horizontal-only and vertical-only weights distributions (Fig 7B insets and 7Ciii). We averaged the mean weight differences between the horizontal and vertical ROIs across the two monkeys to produce the TC of the weight difference in Fig. 7Ci. In Fig. 7Cii we performed a liner fit and computed the Pearson Correlation (r) between the value of weights- difference in the early time (80-100 ms) and the SOA duration, and between the weight difference at the late time (180-200 ms) and the SOA duration. In a similar manner, we calculated separately for each monkey and each SOA the Pearson Correlation (r) between the weight maps obtained for the BM data and the weight maps obtained from the target alone data (e.g. Fig. 5C, 5H; each monkey to its own retinotopic weight-map). Next, we averaged the correlation coefficient across the monkeys to produce the TC of correlation (Fig. 7Di). We then calculated the Pearson Correlation and linear fit with the SOA duration, for the early time and late time (Fig. 7Dii). Finally, we note that we obtained similar results to those obtained by the SVM classifier by using other classifiers e.g. random forest.

### Statistical analysis

Data are presented as mean ± sem (unless indicated otherwise). Nonparametric statistical tests were used: The Wilcoxon rank-sum test to compare between two median of two populations or the Wilcoxon signed-rank test to compare the population’s median to 0. Statistical significance values were corrected for multiple comparisons (Bonferroni correction). In addition, we generated the shuffle condition from the real data as explained in specific parts of the methods. To compute the slope differences for the linear regression of the FGm (Fig. 4F) as function of the SOA as well as the weight difference and correlation to the weight target alone maps as function of the SOA (Fig 7Cii, 7Dii) we used a linear regression model with an interaction term.

## Supporting information

Supplementary material and figures that addressed from the main text

