## Supplementary material and figures that addressed from the main text for "Population responses in V1 encodes stimulus visibility in backward masking"

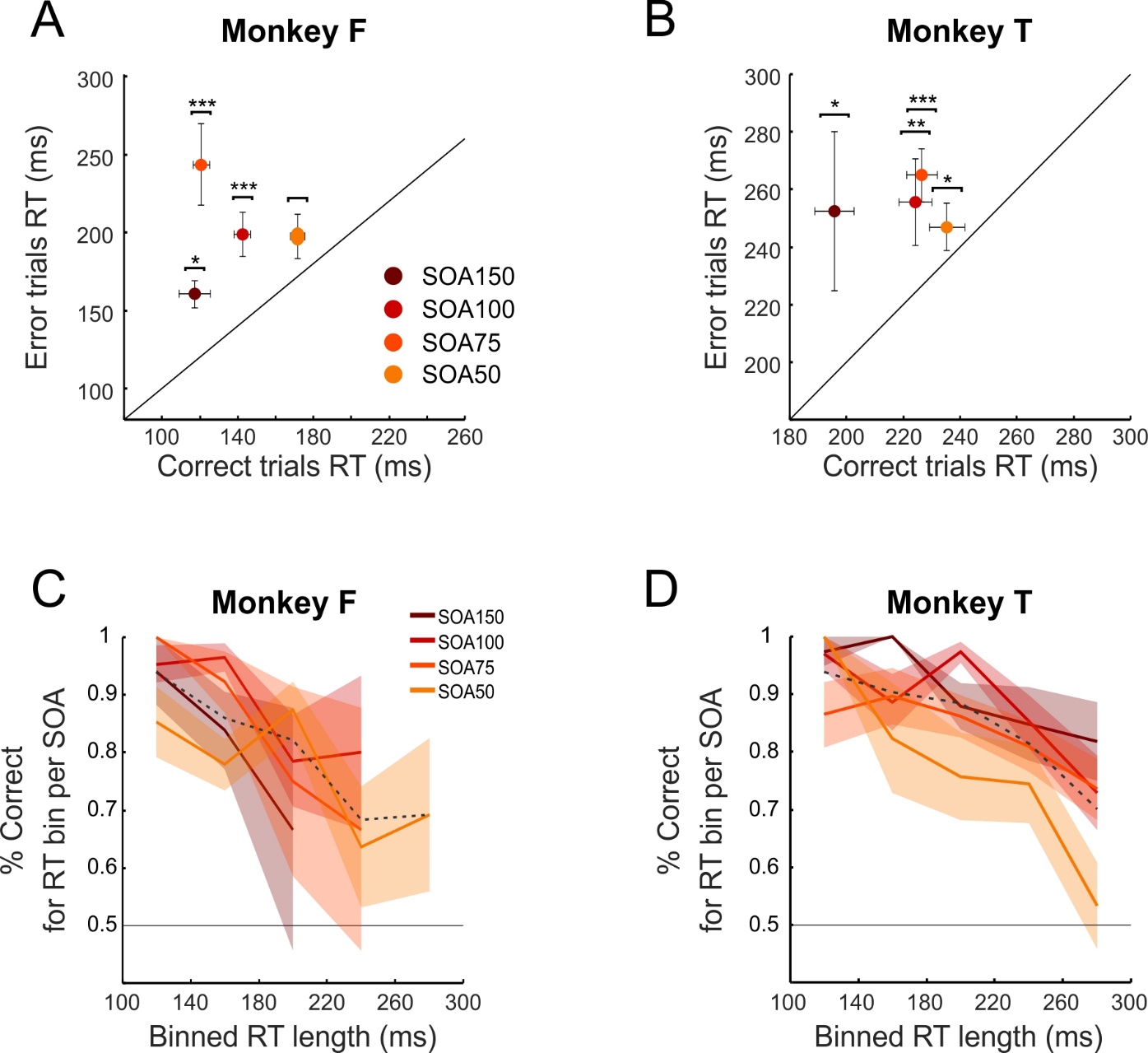

**Supplementary Figure S1 (related to Figure 1). A.** Reaction time of correct trials vs. RT of error trials, data from monkey F. Number of Error trials: n=47, 10, 14 & 9 for SOAs 50, 75, 100 & 150 respectively. Number of Correct trials is as in Fig. 1. Error bars are SEM cross trials, the symbols for p values are as in Fig. 1. **B.** Same as A, but for monkey T. Number of error trials: n=57, 92, 57 & 34 for SOAs 50, 75, 100 & 150 respectively. **C.** Percent of correct trials vs. reaction time, monkey F. The RTs of each SOA were binned into 40 ms duration bins between 100-300 ms. Next, we computed the percent of correct trials for each RT bin, in of an SOA. Dashed gray line represents the mean % correct trials for each bin, when all trials from all SOAs were pooled together. Total number of trials in each SOA: n = 203, 81, 141 & 59 for SOAs 50, 75, 100 & 150 respectively. **D.** Same as C, monkey T. Total number of trials in each SOA: n=143, 310, 263 & 197 for SOAs 50, 75, 100 & 150 respectively.

**Supplementary material 2: Using Convolutional Neural Networks to classify the VSD data (related to Figures 5,6,7).**

To test the validity of the results obtained by the SVM classifier, we analyzed the target alone data and BM data using a different classification approach: Convolutional Neural Networks (CNNs). Unlike SVMs, which treats each pixel as an independent feature, CNNs are better suited for image classification tasks and can capture spatial patterns and spatial correlation between the image pixels (Krizhevsky et al. 2012, LeCun et al. 1998; Goodfellow et al. 2016). In addition, the non-linear activation functions (e.g., ReLU) and pooling layers further enable CNNs to address non-linear relationships and reduce spatial dimensions (LeCun et al. 1998).

**Methods:**

**Data preprocessing:** we used the same data as for the SVM, with one modification. We applied min-max normalization to scale all pixel values between 0 and 1. This step is essential to prevent ReLU activation layers from clipping negative values to zero and to avoid saturation in the sigmoid activation layer.

**Train and test Split and accuracy evaluation**

We used 10-fold cross-validation: in each iteration, 10% of the trials were left out for testing, while the rest were used for training and validation. Data was shuffled prior to the train-test split to minimize the impact of SNR differences between experiments. For the target alone data, an CNN model was trained on the VSD data using a 5 frame window (as in Fig 5) in the rising response time of the VSD signal. For the BM data, we trained separated models for each time window and validated on the 10% sample. The model's accuracy was evaluated using a sliding time window of 5 frames with a stride of 1 frame (as in Fig. 6). To compare the results to the expected by chance, we also trained a separate model on label shuffled data.

**CNN model architecture**

Given the limited dataset size, we focused on shallow CNN architectures to minimize the risk of overfitting. The network architecture comprises three convolutional blocks, each with a convolutional layer followed by a max-pooling layer. The first convolutional layer applies 32 filters of size 3×3 with ReLU activation to input images of 100×100, followed by a 2×2 max-pooling layer. Subsequent convolutional layers utilize 64 filters of size 3×3 with ReLU activation, each followed by a 2×2 max-pooling layer. Post these blocks, a flattening layer converts 2D feature maps into a 1D vector, leading into a dense layer with 64 units and ReLU activation. A dropout layer with a 0.25 rate is then applied. The final layer is a single neuron with a sigmoid activation function for binary classification.

**Features importance**

Unlike SVMs, which rely on explicit feature coefficients, CNNs learn hierarchical patterns through layers of convolution. This makes it challenging to directly interpret the contribution of individual pixels, as is done with linear models. A common approach to address this limitation is to use SHAP (SHapley Additive exPlanations) values or Gradient-weighted Class Activation Mapping (Grad-CAM).

**SHAP**

The SHAP (i.e. SHapley Additive exPlanations) values, based on Shapley values from game theory (Shapley, 1953), offer a framework for interpreting model predictions by attributing contributions to individual features (Lundberg and Lee, 2017). Unlike Grad-CAM (see below), SHAP values show both positive and negative influences on predictions, providing a more granular measure of feature importance. In binary CNN classifiers, SHAP values quantify each pixel's contribution to the model’s prediction (Molnar, 2019). To ensure comparability across models and test sets, we normalized SHAP values by computing their Z-scores, with positive Z-scores indicating above-average contributions to the positive class, and negative Z-scores indicating below-average or negative contributions.

**Grad-CAM**

Gradient-weighted Class Activation Mapping (Grad-CAM) computes the gradients of a target class score with respect to feature maps of a selected convolutional layer, generating a heatmap that highlights regions contributing to a CNN's predictions (Selvaraju et al., 2017). In our binary CNN, the Grad-CAM heatmap identifies regions most influential in predicting the positive class (vertical label). Higher values indicate stronger support, while lower values suggest less influence. To improve resolution, we applied Grad-CAM to the second-to-last convolutional layer, just before the second dropout. Grad-CAM typically highlights positive influences, so for the negative class (horizontal label) we computed Grad-CAM using gradients of the negative class score, thus capturing regions that negatively impact the prediction.

**Supplementary Figure S2 A-B: CNN replicates high accuracy classification of horizontal and vertical target alone trials**

The shaded rectangle in Supplementary Figure S2Ai (monkey F) represents the time window used for model training (same as for SVM). Before stimulus onset, the model's accuracy is near chance level (50%) across all folds. Following the target onset, the model performance rises sharply to 99.7% and remains elevated throughout the trial duration. These results align closely with those obtained using SVMs (Figure 4E). For monkey T, a similar trend is observed, with accuracy peaking at 83% and remaining high above chance level (Supplementary Figure S2Bi). The dashed curve in supplementary Fig. S2Ai and S2Bi shows the accuracy for shuffled data.

Next, we evaluated the predictive importance of individual pixels using SHAP values. Supplementary Figure S2Aii summarizes the grand analysis of mean SHAP values in non-overlapping horizontal and vertical ROIs, as well as in their shared area. For monkey F, the SHAP value is -1.71 ± 0.04 (mean ± SEM) in the horizontal ROI, 1.42 ± 0.033 in the vertical ROI, and 0.04 ± 0.035 in the common ROI. These values are significantly different (p < 0.01, Wilcoxon Rank Sum Test) and are consistent with the SVM-derived pixel weight distributions (Fig. 5). A similar pattern is observed in monkey T (Supplementary Figure S2Bii), with SHAP values of -1.52 ± 0.07 in the horizontal ROI, 1.67 ± 0.06 in the vertical ROI, and -0.137 ± 0.09 in the shared area (p < 0.01, Wilcoxon Rank Sum Test). These findings further confirm consistency between CNN and SVM results and validate the results that were obtained in the SVM for target alone stimulus. In addition, Figures S2Aii inset shows the mean SHAP value maps across all test samples. Warm colors (positive SHAP values) are clustered within the vertical retinotopic ROI, while cold colors (negative SHAP values) are clustered within the horizontal ROI. This spatial distribution highlights the robust contribution of vertical ROI pixels toward the vertical target prediction, and horizontal ROI pixels toward opposing predictions.

**Supplementary Figure S2 C-D: CNN classification of backward masking data confirms accuracy differences between Short and Long SOAs**

Next, we applied CNNs to the BM VSD data. Supplementary Figure S2C depicts the time course (TC) of classification accuracy for the CNN model trained on short SOAs (50 and 75 ms) vs. long SOAs (100 and 150 ms, solid lines). The results are consistent with those obtained using SVMs, showing two distinct peaks of classification accuracy in early and late time (Fig. 6Ai). In the early time, there is almost no difference between long and short SOAs. A significant difference is observed between 110-130ms after stimulus onset (Wilcoxon Rank Sum Test, p value range: < 0.001 to < 0.01). In the late time, the model trained with longer SOAs maintains higher accuracy levels relative to the short SOAs (p < 0.01). Between 220 and 260 ms after onset, its accuracy remains within 0.77± 0.02 to 0.71±0.024, whereas the accuracy with shorter SOAs declines to 0.66± 0.021 to 0.59 ± 0.02. The differences in all time points are significant (p < 0.01). The shuffled data in the BM show chance level throughout the entire time (Fig S2C, dashed curves). Finally, we performed a similar analysis for monkey T which showed smaller differences between long and short SOAs, yet similar trend to that shown in Fig. 6Ci. The difference in model accuracy between long and short SOAs in the late peak was for SOA150: 0.58 ± 0.026 to 0.56± 0.021. vs. SOA 50: 0.53 ± 0.04 to 0.48± 0.046). There was no statistical significance (notably, the stimulus luminance in monkey T was lower, see Methods in main text).

Next, we investigated the feature importance in the CNN model. While CNN lacks a direct equivalent to SVM-derived feature importance, SHAP values provide a means to estimate the relative contribution of each pixel (see Methods). To systematically assess this, we analyzed the mean SHAP values across four conditions: SOA150 in early time, SOA150 in late time, SOA50 in early time, and SOA50 in late time. Supplementary Fig. S2Di shows the mean SHAP weights in the non-overlapping ROIs and in the common ROI for the early and late time for long SOAs (150 and 100). During the early time of the long SOAs, the mean SHAP value in the horizontal ROI is -0.61 ± 0.07 (mean ± SEM), in the vertical ROI 0.06 ± 0.04, and in the common ROI 0.05 ± 0.06 The SHAP value in the horizontal ROI were significantly different from those in the vertical ROI (p < 0.01; Wilcoxon Rank Sum Test). For the late peak of SOA150, the mean SHAP in the horizontal ROI is -0.36 ± 0.21, in the vertical ROI 0.567 ± 0.07 and in the common ROI 0.378 ± 0.1. Again, the SHAP values in the horizontal ROI were significantly different from those in the vertical ROI (p < 0.01; Wilcoxon Rank Sum Test). The same analysis was performed for Short SOAs (75 and 50), while the weights of the SHAP values in the early time, show similar trend to the one observed for the long SOAs early and late time: negative SHAP values in the horizontal ROI (-0.47±0.11^3^, and positive SHAP values in the vertical and common ROIs (0.22±0.036, 0.39±0.098, respectively). In the late time the mean SHAP in the horizontal ROI is positive (0.29±0.95) while vertical and common ROIs maintain the same trend (0.056±0.04, 0.43±0.07, respectively). This shift from negative to positive SHAP values in the horizontal ROI when trained on later frames is aligned with the findings above and provide an additional perspective on the spatial-temporal differences between the two conditions, indicating a temporal evolution of feature importance in the horizontal ROI.

A similar pattern was observed in monkey T, where in long SOA and late time, the mean Z-score SHAP values in the horizontal ROI was negative, while that of the vertical ROI was positive. In contrast, for SOA50 late peak, the mean Z-score SHAP values in horizontal and vertical ROIs were both negative. This provides additional evidence to the lack of spatial organization in the late peak for SOA50, and the difference of the two conditions.

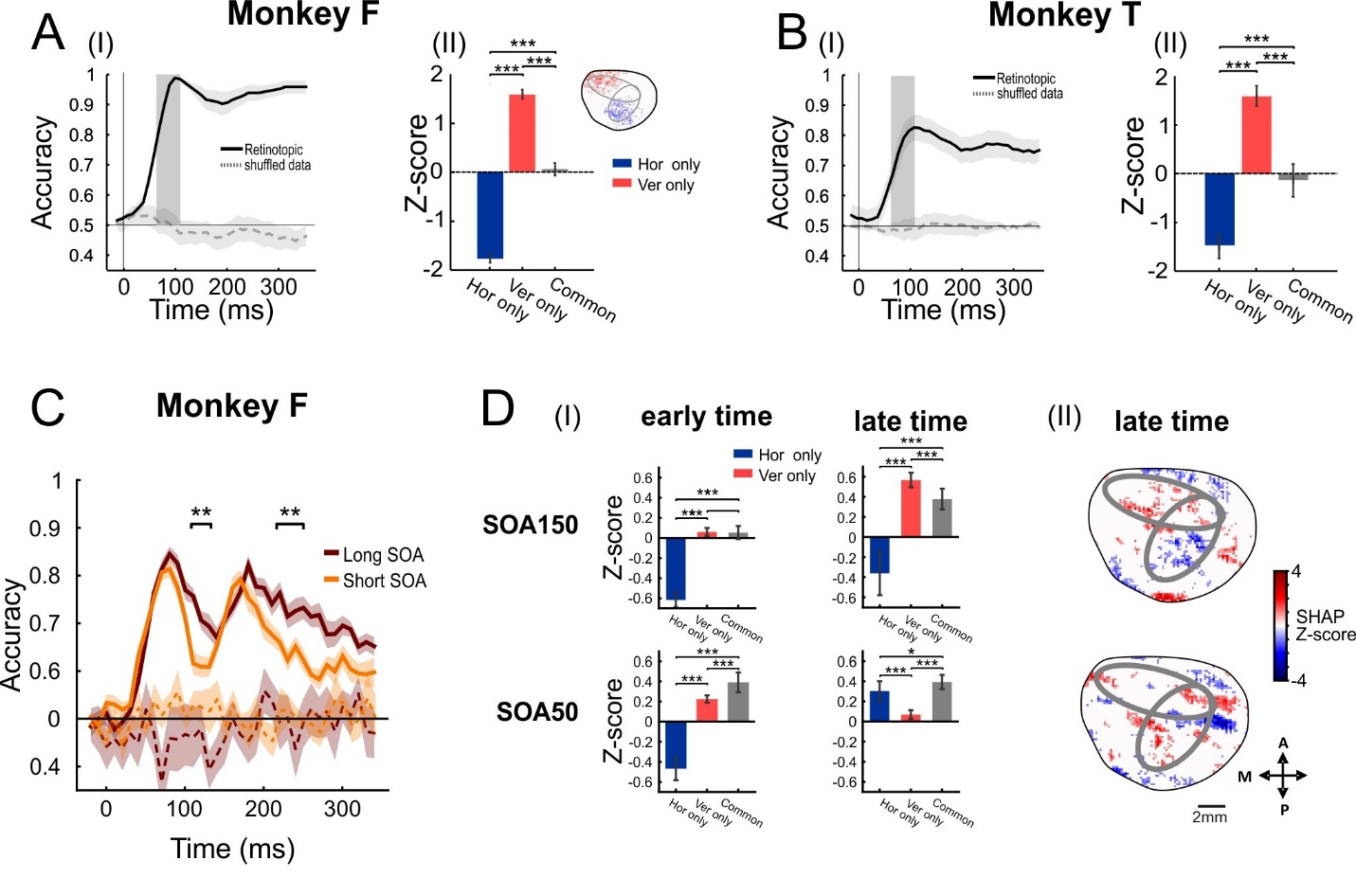
**Supplementary Figure S2: Application of Convolutional Neural Network (CNN) on the VSD data shows similar results to the SVM model. A-B: A CNN classifier on the targets only trials for retinotopic mapping in V1: accuracy and feature importance; C-D: A CNN classifier on the BM trials accuracy and feature importance. A. Right:** Accuracy TC of the CNN classifier. The solid line depicts the classifier accuracy as a function of time, the dashed line represents accuracy for the shuffled data (see Methods). The vertical grey rectangle marks the time window used for model training (70–110 ms after stimulus onset). Shaded areas around the curves indicate SEM across all folds. **Left:** Mean SHAP value at Z-scores units for vertical-only, horizontal-only, and common ROIs. **Top**: Map of SHAP value at Z-score of the classifier, computed as the mean across all folds. Values were Z-scored as described in Methods. For clarity, only the top 10% of the highest and lowest pixel values in V1 are displayed. Red represents pixels that contribute to predicting the positive class (vertical), corresponding to positive SHAP values, while blue represents pixels that increases the likelihood of predicting the negative class (horizontal), corresponding to negative SHAP values, respectively**. B.** Same as A, for monkey T. * p < 0.05; ** p < 0.01; *** p < 0.001 for ranksum test between the ROIs. **C.** Time-course of accuracy for CNN models trained to discriminate between vertical and horizontal trials using a sliding window of 50 ms (see Methods). Each data-point depicts the mean accuracy across all folds. Dashed lines represent the same analysis for the shuffled data. **D(i).** The mean SHAP value at Z-scores units in the vertical-only, horizontal-only, and common ROIs for early and late time-windows (70-90 ms & 170-190 ms after stimulus onset, respectively). **D(ii).** Maps of SHAP value at Z-score for the late time-window (180 ms after stimulus onset), for SOA 50 & 150 (For clarity, only the top 10% highest and lowest values are shown).
